# Genetically Engineered Proportional-Integral Feedback Controllers for Robust Perfect Adaptation in Mammalian Cells

**DOI:** 10.1101/2020.12.06.412304

**Authors:** T. Frei, C.-H. Chang, M. Filo, A. Arampatzis, M. Khammash

## Abstract

To support their survival, cells adapt to environmental disturbances by maintaining a constant internal milieu. Robust perfect adaptation is a strategy that utilizes integral feedback to promote adaptation by robustly driving regulated physiological variables to their pre-disturbance levels. Present in natural systems, this stringent regulatory strategy promises to enable the engineering of sophisticated genetic programs with diverse applications. Here, we present the first synthetic implementations of integral and proportional-integral feedback controllers in mammalian cells. We show that the integral controller robustly and precisely maintains a desired level of a transcription factor, in spite of induced disturbances and network perturbations. Augmenting proportional feedback reduces stochastic variability while maintaining robust perfect adaptation. We demonstrate the benefits of these controllers in mitigating the impact of resource burden and investigate their use in cell therapy. The synthetic biological realization of robust perfect adaptation holds promise for substantial advances in industrial biotechnology and cell-based therapies.

## Introduction

The ability to maintain a steady internal environment in the presence of a changing and uncertain external environment — called homeostasis — is a defining characteristic of living systems [1]. Homeostasis is maintained by various regulatory mechanisms, often in the form of negative feedback loops. The concept of homeostasis is particularly relevant in physiology and medicine, where loss of homeostasis is often attributed to the development of disease [2, 3, 4]. In engineering, the ability of a control system to robustly maintain a controlled dynamical system of interest in a desired state in spite of perturbations is routinely achieved through the use of integral negative feedback. In integral feedback, the deviation of the regulated output of a system from its desired level is measured and mathematically integrated over time and then used to drive the system’s input to counteract the deviations and drive them to zero [5]. A system with integral feedback is known to reject constant disturbances and is also able to perfectly track a desired level commonly referred to as the setpoint. More recently, it has become increasingly evident that integral feedback is a regulatory strategy that drives biological adaptation in several systems [6, 7, 8, 9, 10]. Although integral feedback guarantees robust perfect adaptation, it does not in general prevent large transient deviations. To mediate this, control engineers often augment proportional feedback into their integral feedback control systems. By counteracting such large deviations, proportional-integral feedback also suppresses large stochastic fluctuations around the setpoint and therefore provides tighter regulation than integral feedback alone can achieve [11].

Over the last decade, several experimental studies have constructed genetic systems and cell-based therapies that implement negative feedback to mitigate disease [12, 13, 14, 15]. These, however, rely solely on proportional feedback rather than integral or proportional-integral feedback and are therefore not guaranteed to achieve precise and robust regulation. In 2016, Briat et al. introduced a biomolecular circuit topology that implements integral feedback control for general biomolecular systems [16]. Figure 1(a) depicts an abstract representation of this control motif. A subsequent publication by the same authors showed that additional proportional negative feedback further reduces variance in the controlled output [11]. Central to this strategy — termed antithetic proportional-integral feedback — is the so-called annihilation (or sequestration) reaction between the two species that implement the controller (reaction with rate *η* in Figure 1(a)). The annihilation refers to the requirement that both controller species abolish each other’s function when they interact. Another stringent requirement to achieve integral feedback is that the two controller species on their own remain fairly stable over time. Given these conditions, any network interconnected in a stable way with this antithetic integral controller will achieve robust adaptation (Figure 1(c)). The incorporation of additional proportional negative feedback from the output of the controlled network to the actuation reaction then yields proportional-integral feedback (Figure 1(a)). Independent of integral feedback, this proportional feedback introduces a reduction in the variance of the controlled output (Figure 1(c)).

**Figure 1:**
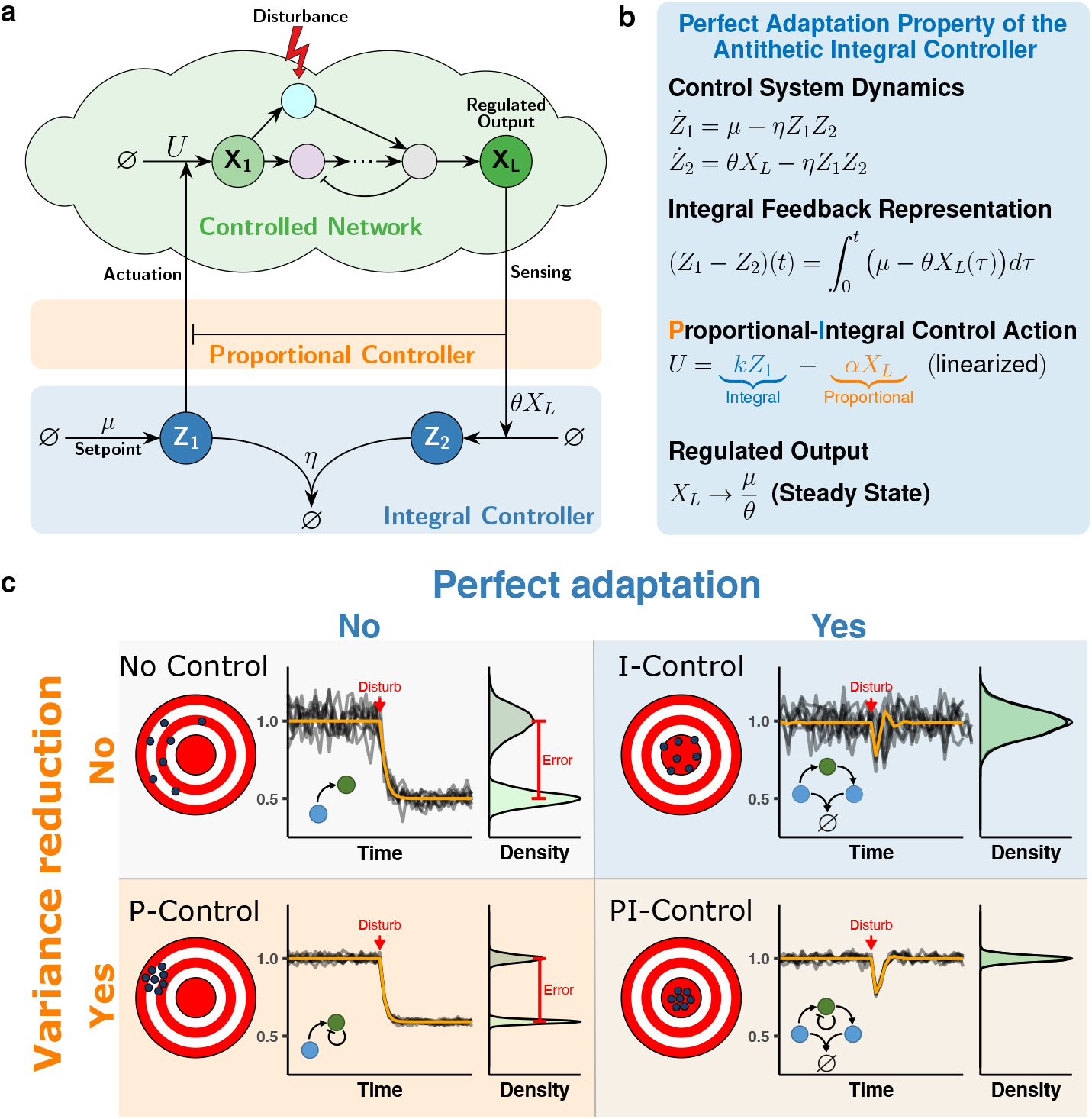
The antithetic proportional-integral feedback motif. **(a)** Network topology of an arbitrary molecular network interacting with an antithetic proportional-integral feedback motif. The nodes labelled with Z_1_ and Z_2_ together compose the antithetic motif responsible for realizing integral feedback. Species Z_1_ is produced at a rate *μ* and is functionally annihilated when it interacts with species Z_2_ at a rate *η*. Furthermore, Z_1_ interacts with the controlled network by promoting the production of species X_1_. To close the feedback loop, species Z_2_ is produced at a reaction rate that is proportional to *θ* and the regulated output species X_L_. An additional negative feedback from the output to the production reaction extends the motif to proportional-integral feedback. **(b) Dynamics of the antithetic integral controller.** Subtracting the differential equations of *Z*_1_ and *Z*_2_ reveals the integral action of the controller that ensures that the steady state of the output converges to a value that is independent of the controlled network parameters. Additionally, through linearization, the individual control actions from the integral and the proportional parts of the antithetic proportional-integral motif can be expressed separately. **(c) The elements of proportional-integral feedback.** Without any feedback control, the output of the controlled network may be highly variable and will likely respond drastically to a disturbance in the network. By adding integral feedback, it can be assured that the output will adapt perfectly to disturbances. Conversely, by adding proportional feedback the variability in the output can be reduced. Combining the two types of feedback reduces the variability of the output while also ensuring perfect adaptation.

The initial theoretical work has motivated the implementation of antithetic integral control in bacteria [17, 18] and in vitro [19]. A quasi-integral controller in ***E. coli*** [20] also relies on a similar topology. In realizing antithetic integral feedback, one of the main challenges is identifying a suitable implementation of the annihilation (or sequestration) reaction [18]. In the bacterial implementation of the antithetic integral feedback motif [18] stable proteins (a *σ* and anti-*σ* factor pair) were used to realize the sequestration reaction. However, this approach is not directly applicable to mammalian cells. Instead, in this work we exploit hybridization of complementary mRNAs to realize this critical reaction. (Figure 2(a)). For the antithetic integral controller to function properly, the sense and antisense RNAs have to be stable such that their degradation is predominantly due to their mutual interaction (via the hybridization reaction). Unlike bacterial RNAs where the majority of mRNAs have half-lives between 3 and 8 minutes [21], mammalian RNAs are much more stable with typical mRNA half lives of several hours [22]. Indeed in human cells, the majority of mRNAs have half-lives between 6 and 18 hours, with an overall mean value of 10 hours [23, 24]. The hybridization of the mammalian sense/antisense RNAs and their stability allow us to realize the antithetic integral controller in mammalian cells. Sense and antisense mRNA have previously been employed to control gene expression in yeast [25] and to build a genetic oscillator in mammalian cells [26]. Furthermore, antisense RNA has shown promise in the treatment of cancer and other genetic diseases as well as infections [27, 28, 29].

**Figure 2:**
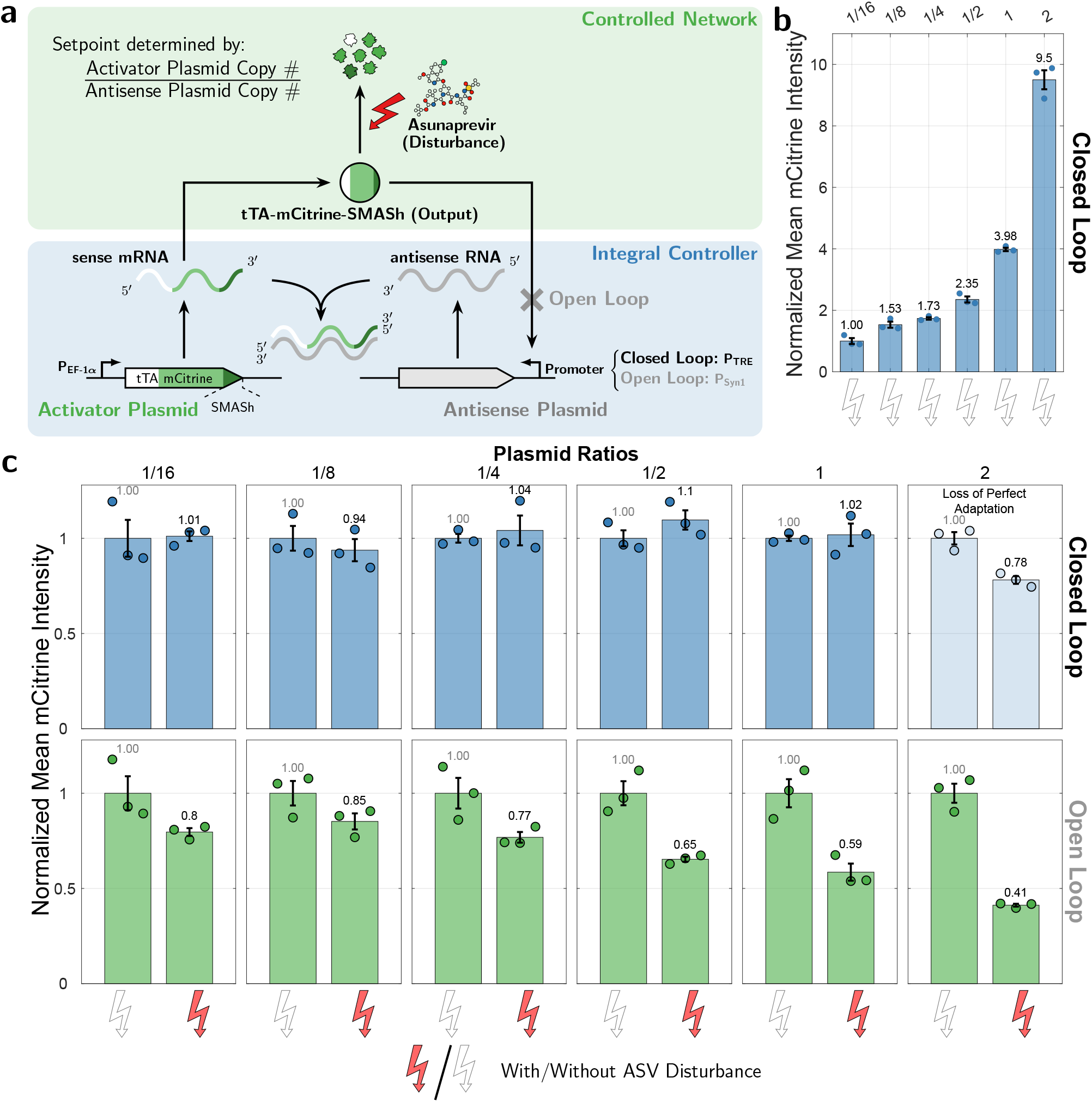
Perfect Adaptation of a Synthetic Antithetic Integral Feedback Circuit in Mammalian Cells. **(a) Genetic implementation of open- and closed-loop circuits.** Both circuits consist of two genes, realized on separate plasmids. The gene in the *activator plasmid* encodes the synthetic transcription factor tTA (tetracycline transactivator) tagged with the fluorescent protein mCitrine and a chemically-inducible degradation tag (SMASh). Its expression is driven by a strong constitutive promoter (P_EF-1 *α*_). The gene in the *antisense plasmid* expresses the antisense RNA under the control of a tTA responsive promoter (P_TRE_). In the open-loop configuration, the TRE promoter was exchanged for a noncognate promoter. In this setting the controlled species is the tTA protein, which can be perturbed externally by addition of Asunaprevir (ASV), the chemical inducer of the SMASh degradation tag. **(b) Steady-state levels of the output (mCitrine) for increasing plasmid ratios.** The genetic implementation of the closed-loop circuit as shown in panel (a) was transiently transfected at different molar ratios (setpoint:= *activator* / *antisense*) by varying the concentration of the activator plasmid while keeping the concentration of the antisense plasmid constant. The data was collected 48 hours after transfection and is shown as mean per condition normalized to the lowest setpoint (1/16) ± standard error for n = 3 replicates. This shows that increasing the plasmid ratio increases the steady-state output level. **(c) Steady-state response of the open-loop and closed-loop implementations to induced degradation by ASV.** The genetic implementation of the open- and closed-loop circuit as shown in panel (a) was transiently transfected at different molar ratios and perturbed with 30 nM of ASV. The data was collected 48 hours after transfection and is shown as mean per condition normalized to the unperturbed conditions for each setpoint separately ± standard error for n = 3 replicates. The unnormalized data is shown in Supplementary Figure A.1 and is provided in a separate file.

Here, we demonstrate perfect adaptation in a sense/antisense mRNA implementation of the antithetic integral feedback circuit in mammalian cells and show that the resulting closed-loop control system is highly robust to network changes and parameter disturbances. By further incorporating proportional feedback to achieve proportional-integral feedback control action, we also increase the tightness of the resulting adaptation. Furthermore, we derive a mathematical (mechanistic) model that describes the various interactions in the system. We show that the obtained model fits the experimentally obtained data well, and is also capable of predicting the robustness features of our implementation of the antithetic integral controller. Lastly, we demonstrate the applicability of our integral and proportional-integral controllers by demonstrating perfect mitigation of gene expression burden and show that the proportional-integral controller provides superior consistency over integral feedback alone.

## Results

A schematic depiction of the sense/antisense RNA implementation of the antithetic integral feedback circuit is shown in Figure 2(a). The basic circuit consists of two genes, which are encoded on separate plasmids. The gene in the *activator plasmid* is the synthetic transcription factor tTA (tetracycline transactivator) [30] fused to the fluorescent protein mCitrine. The expression of this gene is driven by the strong mammalian EF-1 α promoter. This transcription factor drives the expression of the other gene in the *antisense plasmid* via the tTA-responsive TRE promoter. This gene expresses an antisense RNA that is complementary to the *activator* mRNA. The hybridization of these two species realizes the annihilation reaction and closes the negative feedback loop. As an experimental control incapable of producing integral feedback, we built an open-loop analog of the closed-loop circuit, in which the TRE promoter was replaced by a non-cognate promoter. The closed-loop configuration is set up to regulate the expression levels of the *activator* tTA-mCitrine. To introduce specific perturbations to the *activator* we additionally fused an Asunaprevir (ASV) inducible degradation tag (SMASh) to tTA-mCitrine [31].

To show that our genetic implementation of the circuit performs integral feedback we apply constant disturbances with ASV at a concentration of 0.033 μM to HEK293T cells which were transiently transfected with either the open- or the closed-loop circuit. Additionally, we vary the setpoint by transfecting the two plasmids at ratios ranging from 1/16 to 2 (Activator Plasmid/Antisense Plasmid). The fluorescence of the cells was measured 48 hours after transfection using flow cytometry. As the setpoint ratio increases, so does the fluorescence of tTA-mCitrine, indicating that our circuit permits setpoint control (Figure 2(b) and Supplementary Figure A.1). Note that this fluorescence is a monotonically-increasing function of the plasmid ratios (see also the function *θ* in Figure C.7(b)). We consider a circuit to be adapting if its normalized fluorescence intensity stays within 10% of the undisturbed control. Under this criterion, adaptation is achieved for all the setpoints tested below 2 in the closed-loop configuration. In contrast, none of the open-loop configurations manage to meet this adaptation requirement (Figure 2(c)).

Next, we sought to demonstrate that our implementation of the antithetic integral controller will provide disturbance rejection at different setpoints regardless of the network topology it regulates. Therefore, we added a negative feedback loop from tTA-mCitrine to its own production. This negative feedback was realized by the RNA-binding protein L7Ae [32], which is expressed under the control of a tTA-responsive TRE promoter and binds the kink-turn hairpin on the sense mRNA to inhibit translation (Figure 3(a)). The closed- and open-loop circuits were transiently transfected either with or without this negative feedback plasmid to introduce a perturbation to the regulated network. The setpoints 1/4 and 1/2 were tested by transfecting an appropriate ratio of the *activator* to *antisense* plasmids. These different conditions were further perturbed at the molecular level by adding 0.033 μm ASV to induce degradation of tTA-mCitrine. As shown in Figure 3(b) (see also Supplementary Figure A.2), the closed-loop circuit rejects both perturbations nearly perfectly in all cases, whereas again the open-loop circuit fails to adapt.

**Figure 3:**
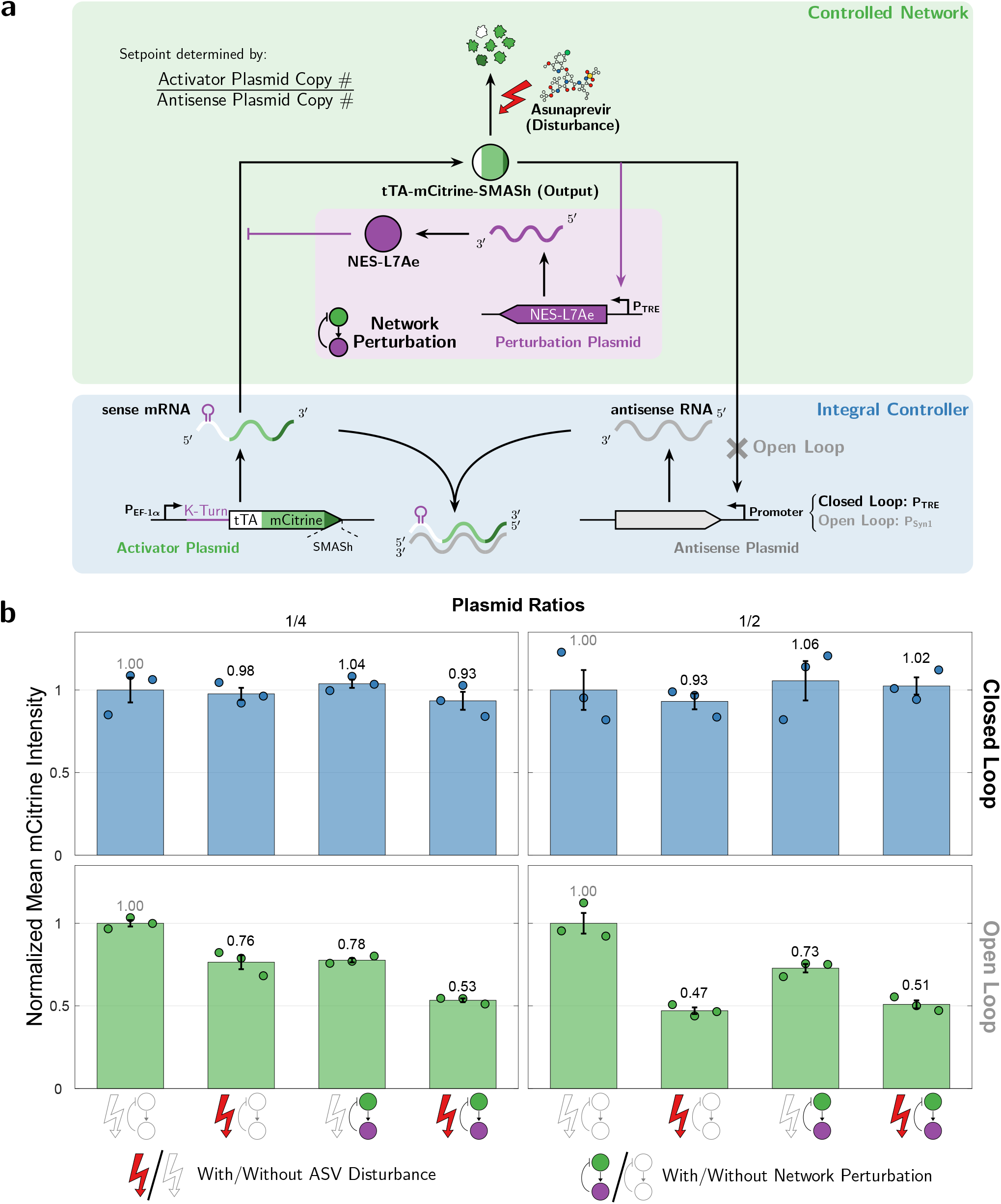
Perturbation to the Controlled Network. **(a) Extension of the network topology with a negative feedback loop.** A negative feedback loop from tTA-mCitrine to its own production was added by expressing the RNA-binding protein L7Ae under the control of a tTA-responsive TRE promoter. This protein binds to the kink-turn hairpin on the sense mRNA to inhibit the translation of tTA. **(b) The closed-loop circuit is not affected by the topology of the regulated network.** The closed- and open-loop circuits were perturbed by co-transfecting the *network perturbation* plasmid and by adding 30 nM of ASV. This was done at two setpoints 1/4 and 1/2 (setpoint:= *activator / antisense*). The HEK293T cells were measured using flow cytometry 48 hours after transfection and the data is shown as mean per condition normalized to the unperturbed network and no ASV condition ± standard error for n = 3 replicates. The unnormalized data is shown in Supplementary Figure A.2 and is provided in a separate file.

The capability of the antithetic integral controller to reject topological network perturbations, as demonstrated previously in Figure 3, allowed us to further improve the controller performance by increasing its complexity. In particular, we implement a common control strategy that is extensively applied in various engineering disciplines, referred to as Proportional-Integral (PI) control. This control strategy adds to the Integral (I) controller Proportional (P) feedback action to enhance dynamic performance, such as transient dynamics and variance reduction [11], while maintaining the adaptation property. To implement proportional feedback control that acts faster than the integral feedback, we use a proxy protein, namely the RNA-binding protein L7Ae, which is produced in parallel with mCitrine-tTA from a single mRNA via the use of P2A self-cleavage peptide (Figure 4a). Therefore, the expression level of L7Ae is expected to proportionally reflect the level of tTA-mCitrine. The negative feedback is hence realized via the proxy protein that inhibits translation by binding the 5’ untranslated region of the sense mRNA. Note that, as opposed to the circuit in Figure 3(a), the production of L7Ae in the PI controller is not regulated by the tTA responsive TRE promoter. Instead, it is directly controlled by the sense mRNA. Furthermore, the proportional feedback realized in the PI controller is expected to act faster than the feedback implemented by the tTA-dependent production of L7Ae (Figure 3) because it does not require additional transcription and translation steps.

**Figure 4:**
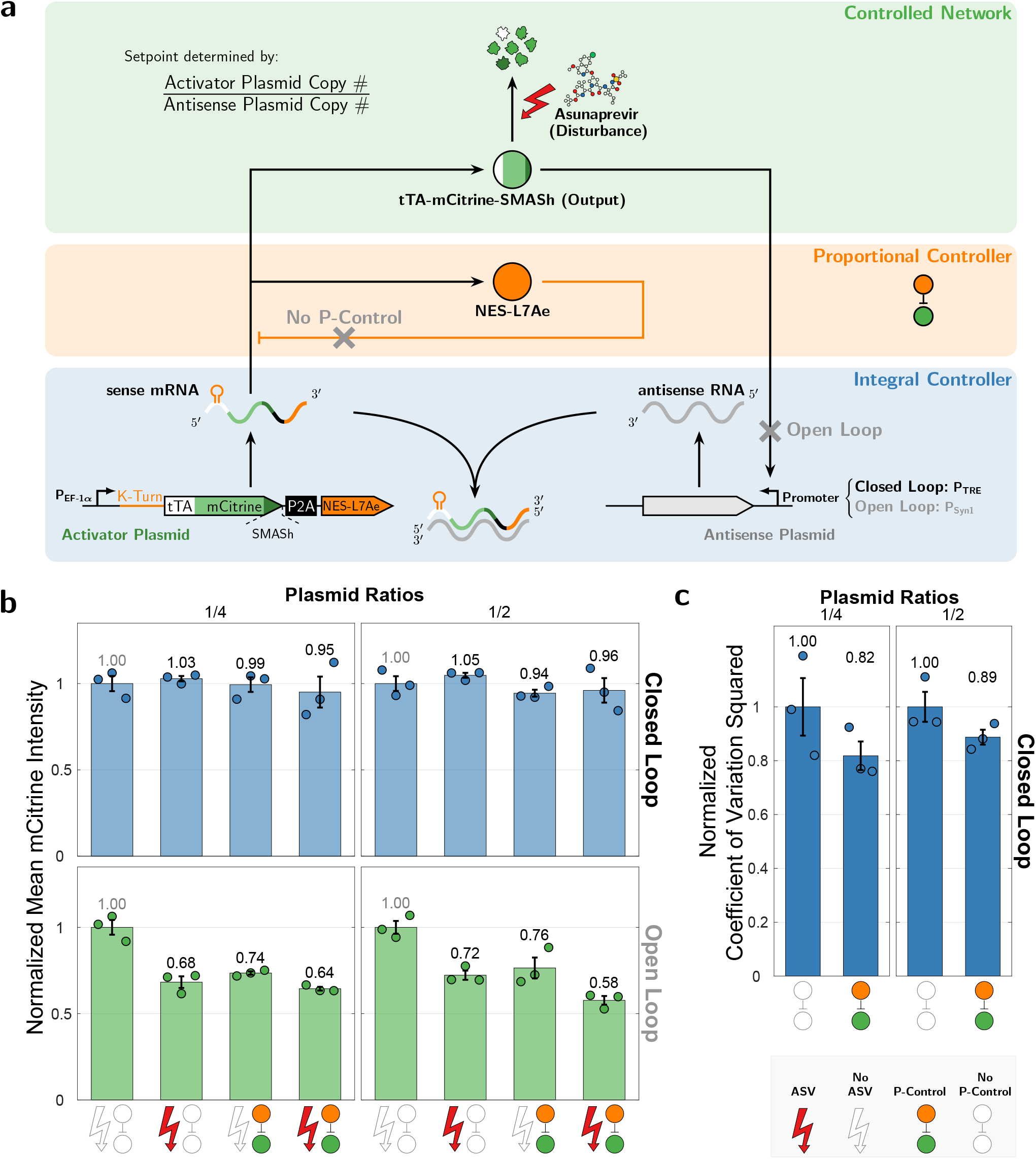
A Proportional-Integral Controller. **(a) Genetic implementation of a Proportional-Integral (PI) controller.** A negative feedback loop from the RNA-binding protein L7Ae (which is proxy to tTA-mCitrine since it is simultaneously produced from the same mRNA) is added to the antithetic motif. This protein binds in the 5’ untranslated region of the sense mRNA species to inhibit the translation of tTA and itself simultaneously. Stronger proportional feedback is realized by adding additional L7Ae binding hairpins. **(b) A PI controller does not break the adaptation property.** The P and PI circuits were implemented by adding a negative feedback loop from L7Ae to the open- and closed-loop circuits. All circuits were perturbed by adding 30 nM of ASV. The HEK293T cells were measured using flow cytometry 48 hours after transfection and the data is shown as mean per condition normalized to the unpertubed (no ASV) condition ± standard error for n = 3 replicates. **(c) Proportional-integral control reduces the steady-state variance.** Computing the normalized coefficient of variation squared on the steady-state flow cytometry distributions, reveals a reduction in variation in the presence of proportional feedback. The coefficients of variation squared were normalized to the No P-Control condition for both setpoints and is shown ± standard error for n = 3 replicates. The unnormalized data is shown in Supplementary Figure A.3 and A.4 and is provided in a separate file.

As illustrated in Figure 4(b), with a standalone Proportional (P) controller, increasing the proportional feedback strength via introducing additional L7Ae binding hairpins has the effect of reducing the steady-state error induced by the drug disturbance. Nonetheless, despite the error reduction, our criteria of adaptation is not met. On the other hand, with a Proportional Integral (PI) controller, the expression of tTA-mCitrine is ensured to be robust to the induced drug disturbance as depicted in Figure 4. This demonstrates that the additional proportional feedback indeed does not break the adaptation property of the antithetic integral controller, as predicted by control theory.

To demonstrate that the three circuits in Figures 2(a), 3(a) and 4(a) are consistent with our understanding of the regulatory topologies, we first derive detailed mechanistic models of these topologies, starting from basic principles of mass-action kinetics. Next, a model reduction technique is carried out based on a quasi-steady-state approximation that exploits the time-scale separation imposed by the various fast binding/unbinding reactions in the network. The mathematical details can be found in Supplementary Information C, D and E, where each circuit is mathematically treated separately. The resulting reduced models are all compactly presented in a single reaction network depicted in Figure 5(a). The overall network can be divided into two biomolecular controller sub-networks – the integral and proportional controllers – that are connected in feedback with another sub-network to be controlled. This is illustrated schematically in Figure 5(a) and mathematically as a set of Ordinary Differential Equations (ODEs) in Figures C.7(b), D.9(b) and E.10(b). The reduced models capture the expression dynamics of the three genes, denoted by **G_1_**, **G_2_** and 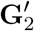, that are encoded in the activator, antisense and network perturbation plasmids, respectively. Gene **G_1_** is constitutively expressed at a rate *μ*(*G*_1_), while the other two genes **G_2_** and 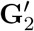 are activated by the (dimer) transcription factor **A** at rates *θ*(*A*; **G**_2_) and 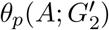, respectively. The derived mathematical expressions of the functions *μ, θ, θ_p_* and the active degradation propensity, λ, are all given in Figures C.7(b) and E.10(b). Note that the model for the circuit of Figure 2(a) (resp. Figure 3(a) can be obtained by setting 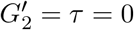 (resp. *τ* = 0); whereas, the model for the circuit of Figure 4(a) can be obtained by setting 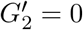 and *τ* = 1.

**Figure 5:**
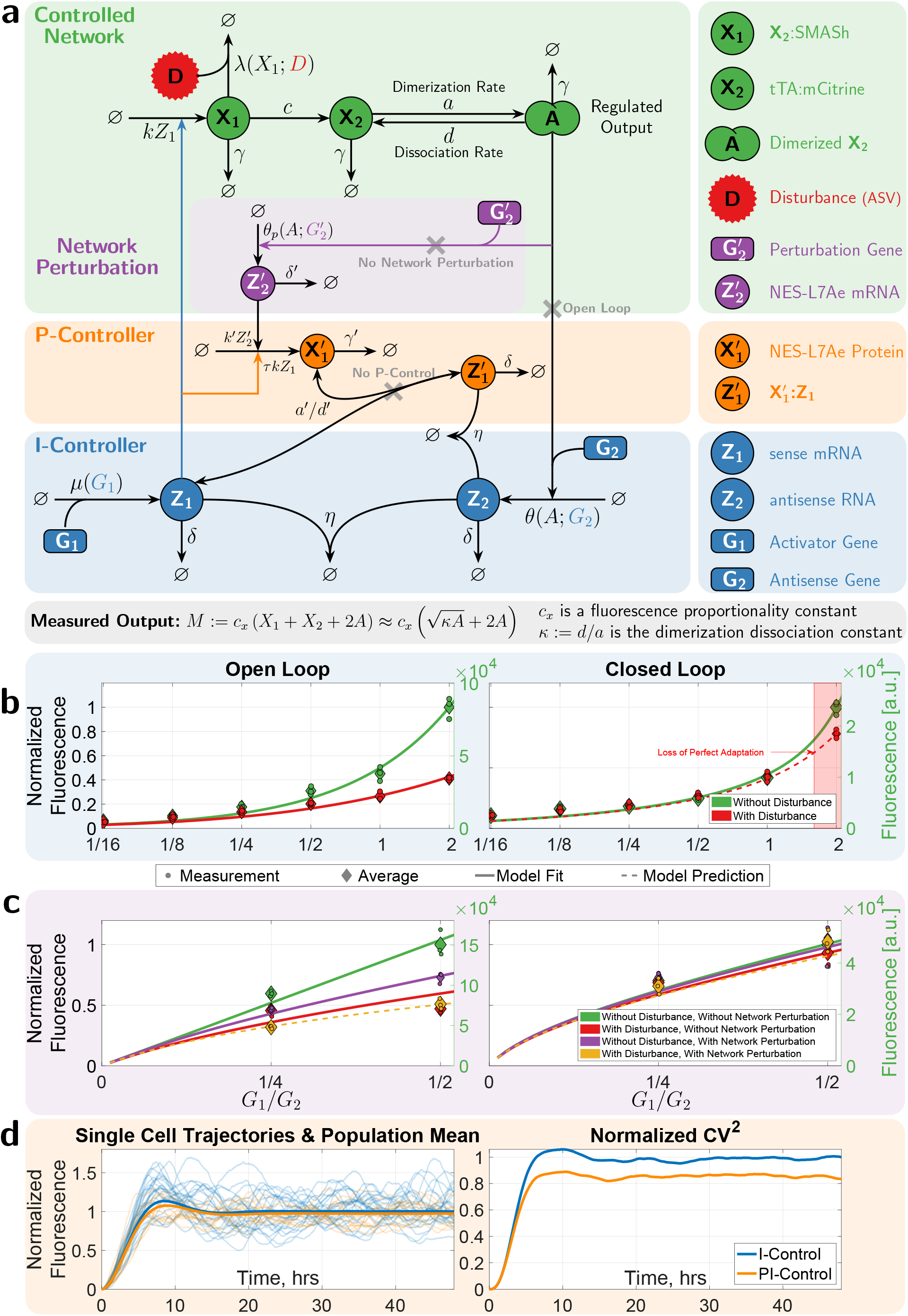
Mathematical Modeling of the Various Circuits. **(a) A Compact Schematic Representation of the Mathematical Models.** This panel describes a chemical reaction network that compactly models the three circuits presented in Figures 2(a), 3(a) and 4(a). The sense mRNA, **Z_1_**, is constitutively produced at a rate *μ*(*G*_1_) that depends on the gene (plasmid) concentration, *G*_1_. Then, **Z_1_** is translated into a fusion of a synthetic transcription factor, fluorescent protein and inducible-degradation tag, referred to as **X_1_**, at a rate *k*. **X_1_** is either actively degraded by the ASV disturbance **D** at a rate *λ*(*X*_1_; *D*) or converted to **X_2_** at a rate *c* by releasing the SMASh tag. The protein **X_2_** dimerizes to form **A** which acts as a transcription factor that activates the transcription of the antisense RNA, **Z_2_**. The transcription rate, denoted by *θ*, is a function of A and the gene concentration *G*_2_. The antithetic integral control, shown in the blue box, is modeled by the sequestration of **Z_1_** and **Z_2_** at a rate *η*. Note that the open-loop circuit is obtained by removing the feedback from the regulated output *A*. The proportional controller, depicted in the orange box, is modeled by producing the protein 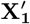, also at a rate *k*, in parallel with **X_1_** to serve as its proxy. A negative feedback is then achieved by the (un)binding reaction between the proxy 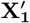 and **Z_1_**. Finally, the network perturbation is modeled via the purple box where an additional gene 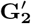 is introduced. This gene is activated by **A** to transcribe the mRNA 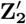 at a rate *θ_p_* which is a function of *A* and 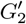. 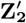 is then translated into the protein 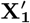 that has, once again, a negative feedback on the production of **X**_1_ by binding to **Z**_1_. See Figures C.7, D.9 and E.10 for a detailed mathematical explanation for each separate circuit. **(b) and (c) Model Calibrations to Experimental Data.** The left plots show the model fits for the open-loop circuits with/without disturbance in (b) and with/without network perturbation in (c). The right plots similarly show the model fits for the closed-loop circuits. The model fits for the open- and closed-loop circuits with/without proportional control are reported in Figure E.10(c). The solid lines denote model fits, while dashed lines denote model predictions. The model fits and predictions show a very good agreement with the experiments over a wide range of plasmid ratios (setpoints) *G*_1_/*G*_2_, with/without disturbance, network perturbation and proportional control. **(d) Stochastic Simulations Demonstrating the Variance Reduction property of the Proportional Controller.** The calibrated steady-state parameter groups of the PI closed-loop circuit, given in (43), are fixed, while the time-related parameters are set as follows: *γ* = *γ′, k* = *c* = *d* = 1 min^−1^ to demonstrate the variance reduction property that is achieved when a proportional controller is appended to the antithetic integral motif. Note that *G*_1_ = 0.002 pmol and *G*_2_ = 0.004 pmol.

Next, we calibrate the derived mathematical models to the experimental measurements that were collected at steady state. The measured fluorescence, denoted by *M*, represents all the molecules involving mCitrine: **X_1_ X_2_**, and **A**. It is shown in Supplementary Information C.4 that M can be expressed solely in terms of the concentration of the regulated output **A**, as shown in the bottom of Figure 5(a), where *c_x_* is an instrument-related proportionality constant that maps concentrations in nm to fluorescence in a.u., and *κ* is the dimerization dissociation constant of **A**. Of course, steady-state measurements alone cannot uniquely estimate all parameters in the model. However, by carrying out a steady-state analysis of the underlying differential equations, we can identify a set of parameter groups (or aggregated parameters) that can be uniquely estimated based on the collected data. The detailed mathematical analyses, showing the aggregated parameter groups and their calibrated values are reported for each circuit separately in Supplementary Information C.5, D.3 and E.3.

In the ideal closed-loop scenario where the dilution/degradation rate *δ* is zero, the steady-state analyses are fairly straight forward and are shown in the bottom of Figures C.7(b), D.9(b) and E.10(b) for each circuit. These analyses show that the steady-state concentration of the regulated output, denoted by *Ā*, is the same for all three circuits and is given by

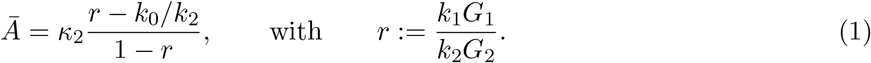

Observe that *Ā* is a monotonically increasing function of the plasmid ratio *G*_1_/*G*_2_, and is independent of the various controlled network parameters, particularly the disturbance *D* and the plasmid concentration 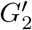. As a result, robust perfect adaptation is exactly achieved since the ASV disturbance and the network perturbation have absolutely no effect on the steady-state concentration of the regulated output **A**.

In practice, the dilution/degradation rate *δ* is never exactly zero, which makes the integrator ‘leaky’. In this case, the steady-state analysis becomes more involved, and one cannot obtain an explicit formula for *Ā* as in the ideal situation. However, implicit (polynomial) formulae can be obtained and are used here to fit the mathematical models to the data. It should be pointed out that when *δ* is sufficiently small relative to other controller rate parameters (as can be achieved with slowly growing cells and fairly stable sense/antisense RNA) the integrator leakiness will be negligibly small, and perfect adaptation can still be achieved for all practical purposes [18, 33]. This is verified experimentally in Figures 2(c), 3(b) and 4(b). The model fits for the integral circuit of Figure 2(a), shown in Figure 5(b), are carried out sequentially for the open-loop circuit first (with and without disturbance), then for the closed-loop circuit (without disturbance). This sequential procedure avoids over-fitting the model to the data. Finally, the closed-loop circuit with disturbance was left for model prediction to assess the calibration accuracy. As shown in the plots of Figure 5(b), the model fits the data very well, and is also capable of predicting the experimentally observed disturbance rejection feature of the antithetic integral controller (dashed red curve in the right plot). Similar model calibration procedures were also carried out for the circuits of Figure D.9(a) and E.10(a), and the model fits and predictions are reported in Figures 5(c) and E.10(c), respectively. Clearly, the models fit the data quite well, and are also capable of predicting another experimentally observed feature of the antithetic integral controller: robustness to network perturbations. The models also show that appending the proportional controller to the integral controller does not affect the steady state of the measured output, but it is capable of reducing the stationary variance (equivalently the coefficient of variation), as demonstrated experimentally in Figure 4(c) (Supplementary Figure A.4(b)) and theoretically through the stochastic simulations depicted in Figure 5(c).

To demonstrate the antithetic integral and proportional-integral controllers in a more practical setting, we apply the circuits introduced in Figure 2 and Figure 4 to decouple the expression of the transcription factor tTA-mCitrine-SMASh from the expression of other genes when they are competing for finite pools of shared resources. This effect was first described in bacteria [34] and later also characterized in mammalian cells [35, 36]. The effective consequence of this is that changes in the expression of one gene inversely affects the expression of all other genes that share a pool of resources with it. In the context of feedback control, the aforementioned changes in gene expression can be seen as disturbances to the controlled network (Figure 6(a)). To experimentally introduce this perturbation, we co-transfected varying amounts of an additional ***disturbance plasmid*** that constitutively expresses the fluorescent protein miRFP670. Previously, it has been observed that the expression of transiently transfected genes is repressed by the presence of double stranded RNA (dsRNA) [37]. We similarly observed that the double stranded RNA (dsRNA) formed through the hybridization of sense and antisense mRNA inhibits the expression of the additionally transfected miRFP670 (comparing Closed Loop to Syn1 Open Loop in Supplementary Figure A.5). To make the gene expression burden — reflected by miRFP670 expression levels — comparable between the closed-loop and open-loop conditions, we replaced the inactive Syn1 promoter with a constitutively active EF1α promoter and tuned the plasmid ratio such that the expression of miRFP670 matches the closed-loop expression (Low EF1α Antisense condition in Supplementary Figure A.5). As was already done in Figure 4, we now compare the responses of the open-loop (No Control), proportional feedback (P-Control), integral feedback (I-Control) and proportional-integral (PI-Control) variant to this new disturbance. As can be seen in Figure 6(b) (Supplementary Figure A.6(a)), a setpoint of 1/2 is maintained within 10 % up to a disturbance strength of 2.3 for I-Control and for all disturbance strengths for PI-Control (Figure 6(b) and Supplementary Figure A.6(a)). This is not the case for the No Control and P-Control configurations, where the steady-state error steadily increases with the increasing strength of the disturbance (Figure 6(b) and Supplementary Figure A.6(a)). In all cases, the disturbance is similar in relative extent (Figure 6(b) top and Supplementary Figure A.6(a) top). In addition to providing perfect adaptation, PI-Control improves regulation over I-control by further reducing the steady-state cell-to-cell variability (Figure 6(c) and Supplementary Figure A.6(b)).

**Figure 6:**
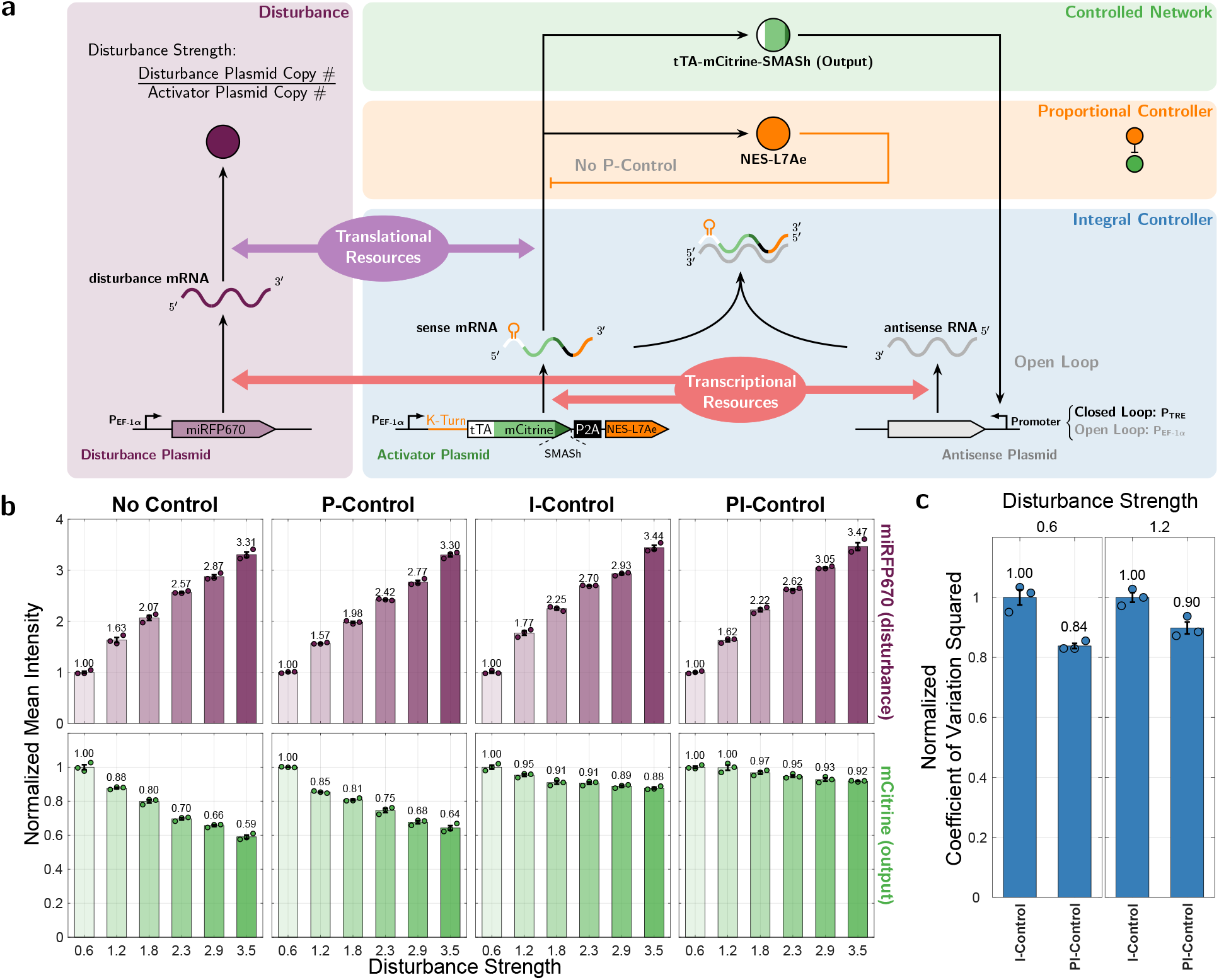
Mitigating Competition for Shared Limited Resources with Antithetic Integral and Proportional-Integral Feedback. **(a) A genetic implementation of an antithetic integral and proportional-integral feedback circuit for mitigating the effects of limited shared resources.** The antithetic integral and proportional-integral feedback circuit characterized in Figure 2 and Figure 4 are re-purposed to mitigate the coupling of gene expression induced by shared pools of finite resources. Varying the amounts of an additional *disturbance plasmid* that constitutively expresses the fluorescent protein miRFP670 introduces a disturbance to the amount of available resources which indirectly affects the expression levels of tTA-mCitrine-SMASh. **(b) Steady-state rejection of disturbances to available limited shared resources.** The *activator plasmid* and *antisense plasmid* for all conditions were transiently transfected at a setpoint ratio of 1/2 together with disturbance strengths varying from 0.6 to 3.5. The disturbance strength describes the amount of disturbance plasmid relative to the activator plasmid. **(c) Reduction in cell-to-cell variability as a result of proportional-integral feedback control.** The coefficient of variation squared was computed for the first two disturbance strengths and normalized two the I-Control condition. The data is shown as the mean ± standard error for *n* = 3 replicates per condition. The unnormalized data is shown in Supplementary Figure A.6 and is provided in a separate file.

## Discussion

This study presents the first implementation of integral and proportional-integral feedback in mammalian cells. With our proof-of-principle circuit we lay the foundation for robust and predictable control systems engineering in mammalian biology. We believe proportional-integral feedback systems will have a transformative effect on the field of synthetic biology just like they have had on other engineering disciplines.

Based on the antithetic motif (Figure 1(a)), we designed and built a proof-of-concept circuit capable of perfect adaptation. This was achieved by exploiting the hybridization of mRNA molecules to complementary antisense RNAs. The resulting inhibition of translation realized the central sequestration mechanism. Specifically, we expressed an antisense RNA through a promoter that was activated by the transcription factor tTA. This antisense RNA was complementary to and bound with the mRNA of tTA to close the negative feedback loop (Figure 2(a)). We further highlighted the properties of integral feedback control by showing that our circuit permits different setpoints. By applying a disturbance to the regulated species we showed that the closed-loop circuit achieved adaptation and provided superior robustness compared to an analogous open-loop circuit (2(c)). Further, we showed that adaptation was also achieved when the setpoint of the circuit was changed. An earlier implementation of the antithetic integral feedback motif in bacteria [18] used a *σ* and anti-*σ* factor pair to realize the sequestration reaction. Due to the requirement of factors native to the bacterial cell for σ factors to activate transcription, this approach is not directly applicable to mammalian cells. Conversely, the sense and antisense RNA approach utilized in this study is likely to be more difficult to realize in bacterial cells due to rapid mRNA turnover.

Moreover, we demonstrated that our realization of the antithetic integral feedback motif is agnostic to the network structure of the regulated species. This was achieved by introducing a perturbation to the controlled network itself (Figure 3(b) and Supplementary Figure A.2). Furthermore, we also demonstrated that the closed-loop circuit still rejected disturbances even in the presence of this extra perturbation to the network. In the open-loop circuit, the disturbance, perturbation, and perturbation with disturbance all led to a strong decrease in tTA-mCitrine expression.

Next, we used the perturbation to the controlled network to incorporate proportional feedback into our integral control circuit directly. We then showed that this proportional-integral feedback controller maintained the same setpoint as the integral controller, even when challenged with induced degradation of the controlled species. To demonstrate that this new controller did utilize proportional feedback, we showed a reduction in the cell-to-cell variability by computing the coefficient of variation squared on the measured fluorescence distributions.

To test our understanding of the mechanistic interactions within our circuits, we derived mechanistic mathematical models for the circuits, starting from basic mass-action kinetics, and showed that the obtained models were capable of fitting the experimental measurements. We also showed that the models were capable of predicting key features of our implementation of the antithetic proportional-integral controller: disturbance rejection and robustness to network perturbations.

Finally, we employed our integral and antithetic-integral feedback circuits to perfectly mitigate gene expression burden on the controlled species caused by introducing an additional, constructively expressed fluorescent protein at varying levels. In light of recent studies on the effects of shared cellular resources in mammalian cells [35, 36], it is important to point out that the dependence of the production of the two controller species on the same resource pool (e.g. transcriptional resources for sense/antisense RNAs) was crucial for maintaining the setpoint despite variations in resource availability. This derives from the fact that the setpoint is a function of the ratio of the production rates of the two controller species (ratio r in 1). Whenever both rates depend similarly on the same resource pool, the effect of this dependence cancels out. When the production rates depend on different resource pools, they do not cancel out and the setpoint becomes sensitive to resource allocation.

Aside from realizing integral feedback control, the sense and antisense RNA implementation is very simple to adapt and is versatile. Indeed both sense and antisense are fully programmable, with the only requirement that they share sufficient sequence homology to hybridize and inhibit translation. Due to this fact, mRNAs of endogenous transcription factors may easily be converted into the antithetic motif simply by expressing their antisense RNA from a promoter activated by the transcription factor. However, one should note that in this case the setpoint to the transcription factor will be lower than without the antisense RNA due to the negative feedback. Furthermore, if the mRNA of the endogenous transcription factor is not very stable, the integrator is expected to not perform perfectly.

Genetically engineered controllers have desirable properties as treatment strategies for homeostasis-related pathologies. Previously, it has been demonstrated that when encapsulated insulin-producing designer cells were implanted in diabetic mice they alleviated the effects of type 1 diabetes mellitus (T1DM) by secreting insulin in response to low blood pH mediated by diabetic ketoacidosis [41] or, alternatively, in response to sensed glucose [15]. This pioneering work provided a proof-of-concept for the practical feasibility of this approach. In this previous work, however, the designed feedback controller is similar to standalone proportional controller, and therefore cannot exhibit the property of robust perfect adaptation that is characteristic of integral feedback. We next exploit our antithetic proportional-integral controller implementation to carry out a simulation study that demonstrates the achievable robust precision and accuracy of the glucose response in modeled diabetic patients. To illustrate the clinical translatability of our proposed controller topologies, we employed disease models for diabetes mellitus (DM) and interfaced them with the different controller circuits (Figure 7). The ability of pancreatic *β*-cells to synthesize and release insulin determines the classification of DM into two main categories: type 1 DM of autoimmune etiology and type 2 DM (T2DM). As a result, we utilized mathematical models for both T1DM [42, 43] and T2DM [38], which originated from the UVA-PADOVA Type 1 Diabetes Simulator (S2008) and its updated versions [42, 44]. This simulator constitutes the first computer model approved by the FDA as an alternative to preclinical trials and animal testing.

**Figure 7:**
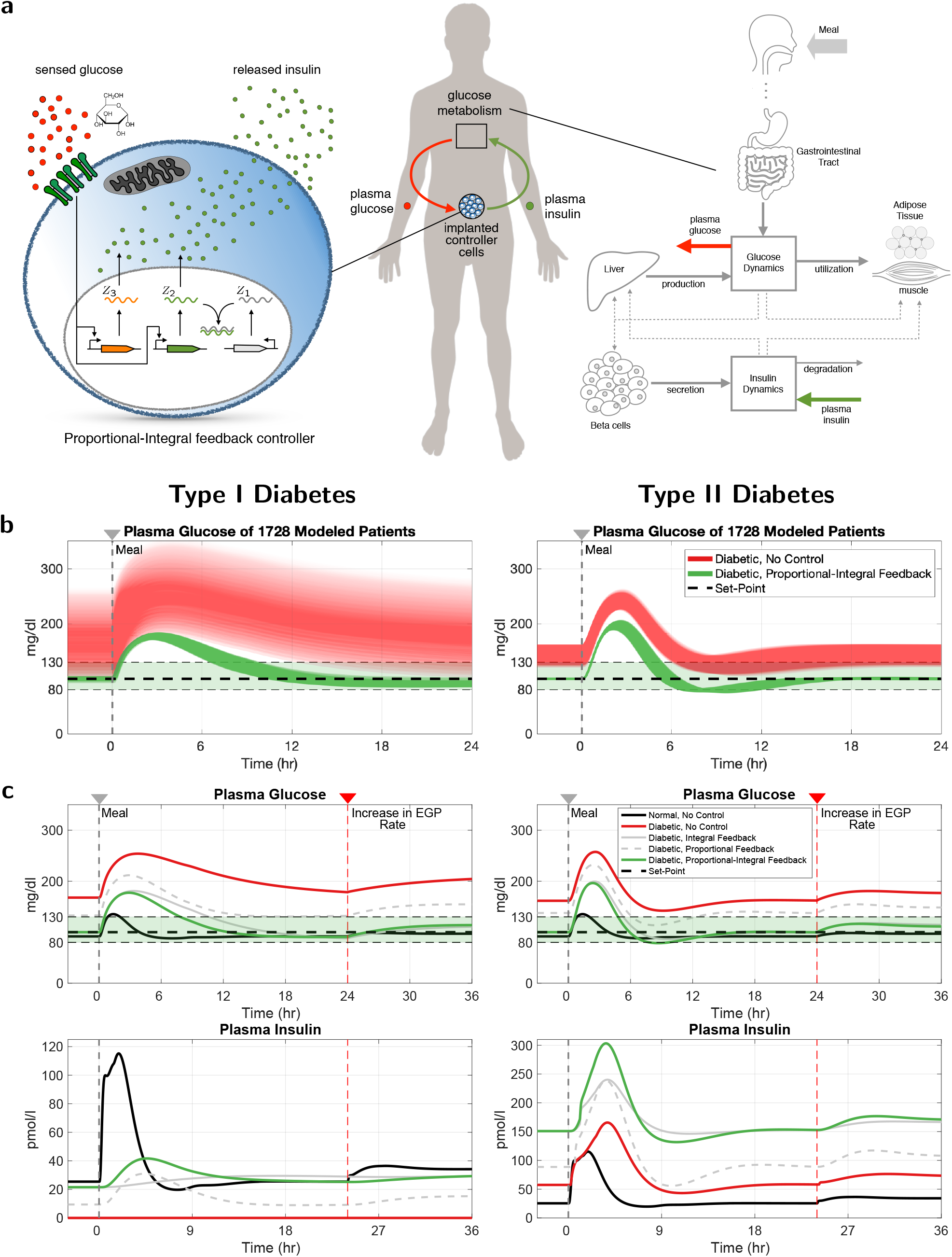
Simulation of Glucose Regulation in the Blood with Antithetic Proportional-Integral Control. **(a) A schematic representation describing the mathematical model of the closed-loop network.** The diagram to the right provides a high-level description of the modeled glucose and insulin dynamics based on [38]. This diagram represents the controlled network, where the output of interest (to be controlled) is the glucose concentration (mg/dl) in the plasma; whereas, the input that actuates this network is the insulin concentration (pmol/l) in the plasma. Note that, unlike the controlled network in the previous figures, this network has a negative gain: increasing the input (insulin) decreases the output (glucose). Hence, to ensure an overall negative feedback, a P-Type controller (with positive gain) is adopted here and shown in the schematic to the left which models a genetically embedded antithetic proportional-integral controller. The P-Type property of the integrator is achieved by switching **Z_1_** with **Z_2_**, that is, the antisense RNA is now constitutively produced while the sense mRNA senses the output (glucose) and actuates the input (insulin). The P-Type property of the proportional controller is achieved by using an activation reaction (instead of an inhibition reaction as in Figure 4(a)) where glucose activates a gene (in orange) to produce insulin. **(b) Robustness to inter-patient variability.** To demonstrate the robustness of our proportional-integral controllers, three parameters *k*_*p*_1__ ∈ [2.4, 3], *V_mx_* ∈ [0.024, 0.071] and *k*_*e*_1__ ∈ [0.0003, 0.0008] (see [38]) in the controlled network are varied, while the controller parameters are fixed. Changes of *k*_*p*_1__ depict alterations in endogenous glucose production (e.g. in various catabolic or stress states [39]), *V_mx_* is used to simulate variations in the insulin-dependent glucose utilization (*U_id_* in [38]) in the peripheral tissues (e.g. by physiological or pathological changes in GLUT4 translocation), while *k*_*e*_1__ is the glomerular filtration rate. The responses are shown for a meal of 40g of glucose at *t* = 0. Adaptation is achieved for all these parameters and for both Type I and II diabetic subjects. **(c) Response to** 40 g **of glucose at time** *t* = 0 **and a disturbance in endogenous glucose production (EGP) rate at** *t* = 24hr. A single meal comprised of 40 g of glucose and an increase of endogenous glucose production rate from *k*_*p*_1__ = 2.7mg/min → 3mg/min (see [38]) are applied to the models of healthy and diabetic subjects at *t* = 0 and *t* = 24hr, respectively. The top (resp. bottom) plots depicts the response of glucose (resp. insulin) concentration; whereas the left (resp. right) plots correspond to a Type I (resp. II) diabetic subject. The black curves correspond to a healthy subject whose glucose levels quickly returns back to the glycemic target range (for adults with diabetes) [80, 130] mg/dl [40] after the meal due the naturally secreted insulin. In contrast, the red curves correspond to uncontrolled diabetic patients whose glucose levels are incapable of returning back to the healthy range due to lack of insulin (Type I) or low insulin sensitivity (Type II). Finally, the solid gray, dashed gray and green curves correspond to diabetic patients whose glucose levels are controlled by our integral, proportional and proportional-integral controllers, respectively. Both integral and proportional-integral controllers are capable of restoring a healthy level of glucose concentration by tuning the set-point to a desired value (100mg/dL); whereas, the proportional controller alone is neither capable of returning to the desired set-point nor rejecting the disturbance. Furthermore, the proportional-integral controller outperforms the standalone integral controller by speeding up the convergence to the set-point, especially for type I diabetes.

Over the past decades the prevalence of DM has increased exponentially, and DM is now considered the most common endocrine disease, affecting approximately 1 in 11 adults globally [45]. Our results propose a closed-loop alternative to open-loop replacement therapy with exogenous insulin, which in the case of T1DM is prescribed for life. They also offer a potentially more manageable approach to the combination of lifestyle changes and pharmacological interventions that is recommended for addressing T2DM management [46]. Moreover, we showed that the simulated glucose control is robust to interpatient variability (see Figure 7(b)), for example due to differences in endogenous glucose production by the liver (clinically found under stress conditions or in critically ill patients [39]), or to changes in renal function, such as physiological or pathological (e.g. diuretic administration, chronic kidney disease) variations in glomerular filtration rate. It was also shown in Figure 7(c) that the antithetic integral and proportional-integral controllers were capable of achieving robust adaptation. In contrast, a standalone proportional controller did not meet the desired set-point, nor could it reject disturbances such as an increase in endogenous glucose production rate (*k*_*p*_1__ in [38]). Note that dissimilarities in the response of the healthy patient and that of the PI-controller-treated patient are, for the most part, not due to any differences between the two regulation strategies (natural vs synthetic). Rather they are mostly attributed to the fact that for the treated patient the insulin was modeled to be synthesized de *novo* from a genetically engineered synthetic insulin gene, leading to inevitable gene expression delay. In comparison, for healthy patients insulin is stored in vesicles for quick release, which ensures a more rapid response — a fact that was also accounted for in the model of the healthy patient. Nevertheless, the response of the PI-controller-treated patient in Figure 7(c) meets all the preprandial and peak postprandial plasma glucose guidelines of the American Diabetes Association [40], and hence offers a potentially effective treatment strategy. Interestingly, the same controller for the single T1DM patient of Figure 7(c) (left) was capable of meeting the guidelines for *all* 1728 patients in Figure 7(b) (left) without requiring re-tuning for different patients — a clear demonstration of robust adaptation. A similar robust adaptation was seen in T2DM, where a single controller met the guidelines for the majority of patients. For those patients for whom the guidelines were not met, the violation was slight (glucose levels exceeded 180 mg/dl only briefly beyond the maximum of two hours Figure 7(b) (right)). This, however, can be remedied by slightly re-tuning the controller for these patients if necessary. The details of the mathematical modeling can be found in Supplementary Information F.

We believe that the ability to precisely and robustly regulate gene expression in mammalian cells will find many applications in industrial biotechnology and biomedicine. In the area of biomedicine, these robust perfectly adapting controllers can be used to restore homeostasis in the treatment of metabolic diseases, as well as for applications in immunotherapy and precise drug delivery.

## Methods

### Plasmid construction

Plasmids for transfection were constructed using a mammalian adaptation of the modular cloning (MoClo) yeast toolkit standard [47]. Custom parts for the toolkit were generated by PCR amplification (Phusion Flash High-Fidelity PCR Master Mix; Thermo Scientific) and assembled into toolkit vectors via golden gate assembly [48]. All enzymes used for applying the MoClo procedure were obtained from New England Biolabs (NEB).

### Cell culture

HEK293T cells (ATCC, strain number CRL-3216) were cultured in Dulbecco’s modified Eagle’s medium (DMEM; Gibco) supplemented with 10 % FBS (Sigma-Aldrich), 1x GlutaMAX (Gibco) and 1 mm Sodium Pyruvate (Gibco). The cells were maintained at 37 °C and 5 % CO_2_. Every 2 to 3 days the cells were passaged into a fresh T25 flask. When required, surplus cells were plated for transfection.

### Transfection

Cells used in transfection experiments were plated in a 96 wells plate at 10,000–15,000 cells per well or in a 24 wells plate at 70,000 - 80,000 cells per well approximately 24 h before treatment with the transfection solution. Alternatively, for the experiments shown in Figure 6, the cell suspension diluted to approximately 140,000 - 160,000 cells per well was used to quench the transfection solution directly. 24-well plates were then seeded from the resulting solution. The transfection solution was prepared using Polyethylenimine (PEI) MAX (MW 40000; Polysciences, Inc.) at a 1:3 (μg DNA to μg PEI) ratio with a total of 100 ng plasmid DNA for the 96 wells plate or 500 ng plasmid DNA for the 24 wells plate. The specific amounts of plasmid and cells used for each experiment are summarized in Supplementary Tables B.1, B.2, B.3, B.4, B.5 and B.6. All plasmids used for transfection are summarized in Supplementary Table B.7. The solution was prepared in Opti-MEM I (Gibco) and incubated for approximately 25 min prior to addition to the cells.

### Flow cytometry

Approximately 48 h after transfection the cells were collected in 60 μL Accutase solution (Sigma-Aldrich). The fluorescence was measured on a Beckman Coulter CytoFLEX S flow cytometer using the 488 nm laser with a 525/40+OD1 bandpass filter. For each sample the whole cell suspension was collected. In each measurement additional unstained and single color (mCitrine only) controls were collected for gating and compensation.

## Data analysis

The acquired data was analyzed using a custom analysis pipeline implemented in the R programming language. The measured events are automatically gated and compensated for further plotting and analysis.

## Acknowledgments

We thank Dr. Gabriele Lillacci and Eline Bijman for their support during the early stage of the project. We also thank Drs. Stephanie Aoki and Ankit Gupta for reading the manuscript and providing many useful comments. M.K. acknowledges funding from the European Research Council (ERC) under the European Union’s Horizon 2020 research and innovation programme (CyberGenetics; grant agreement 743269).

## A Supplementary Figures

**Figure A.1:**
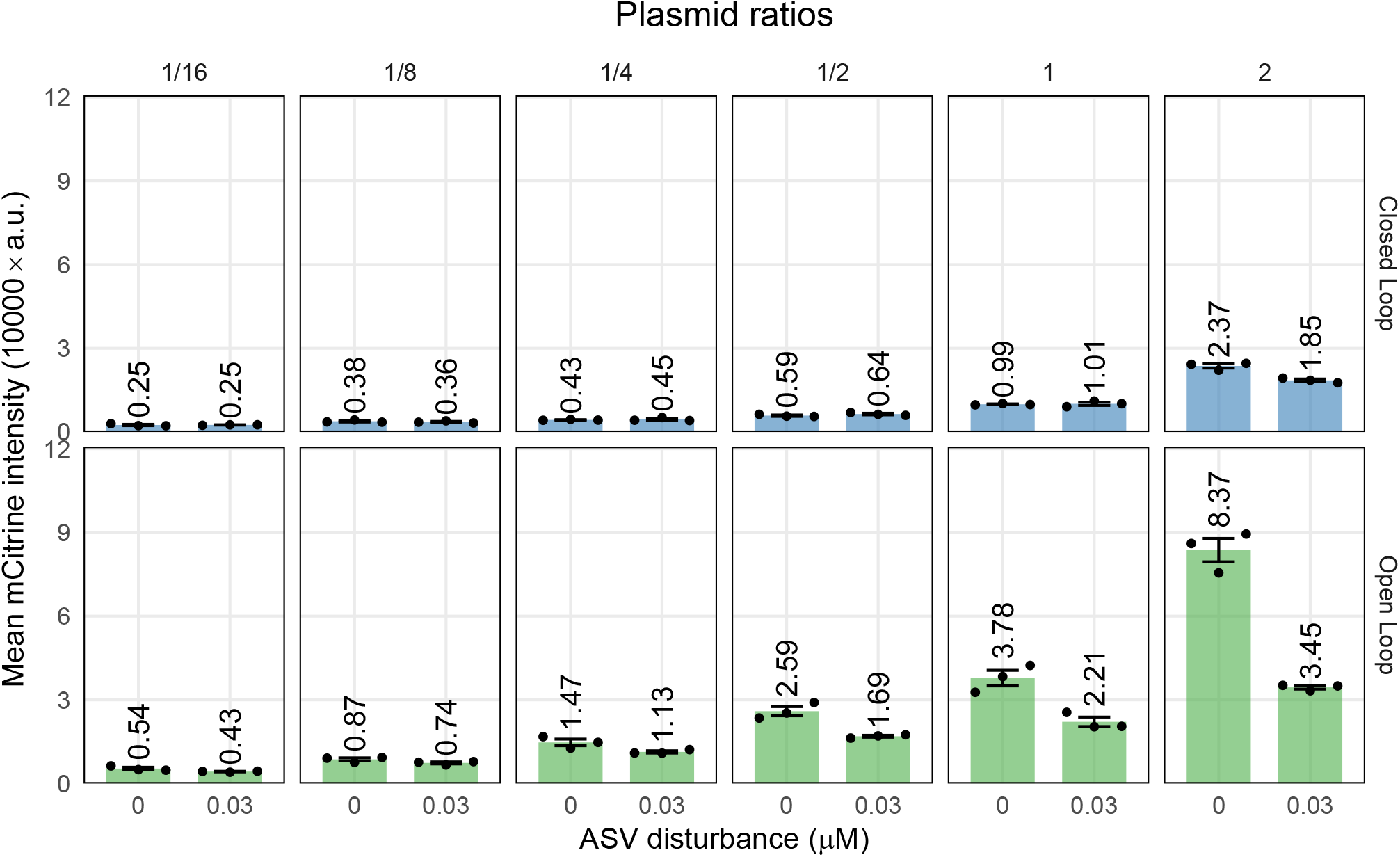
Fluorescence values shown in Figure 2 in arbitrary fluorescence units. The genetic implementations of the open- and closed-loop circuits as shown in Figure 2(a) were transiently transfected in six different molar ratios (setpoint:= activator/antisense) and perturbed with 30 nM ASV. The data was collected 48 hours after transfection and is plotted as mean fluorescence intensity ± standard error for N = 3 replicates. The data is provided in a separate file.

**Figure A.2:**
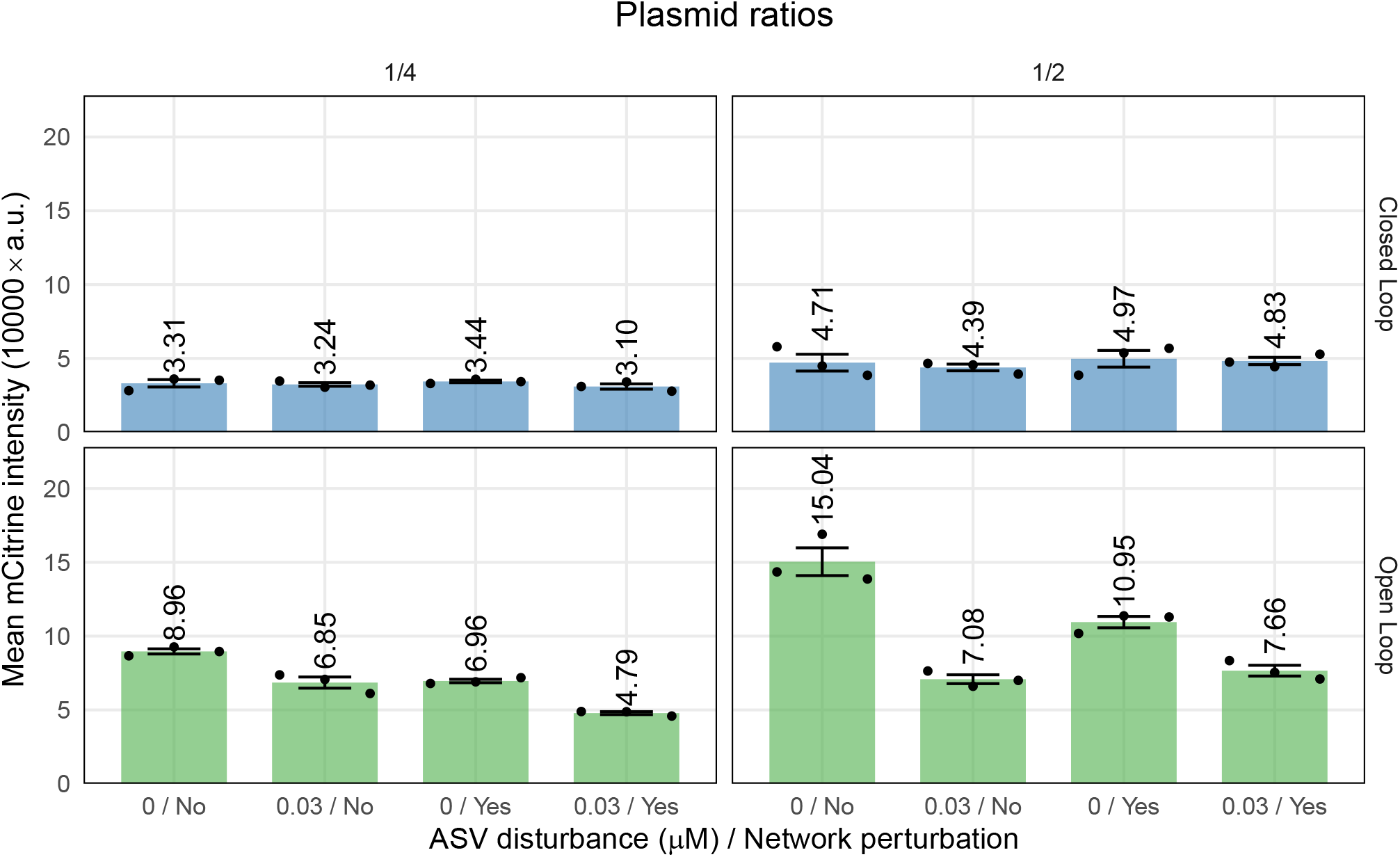
Fluorescence values shown in Figure 3 in arbitrary fluorescence units. The closed- and open-loop circuits were perturbed by an additional negative feedback loop from L7Ae and by adding 30 nM ASV, as shown in Figure 3(a). This was done for two setpoints 1/4 and 1/2 (setpoint:=activator/antisense). The data was collected 48 hours after transfection and is plotted as mean fluorescence intensity ± standard error for N = 3 replicates. The data is provided in a separate file.

**Figure A.3:**
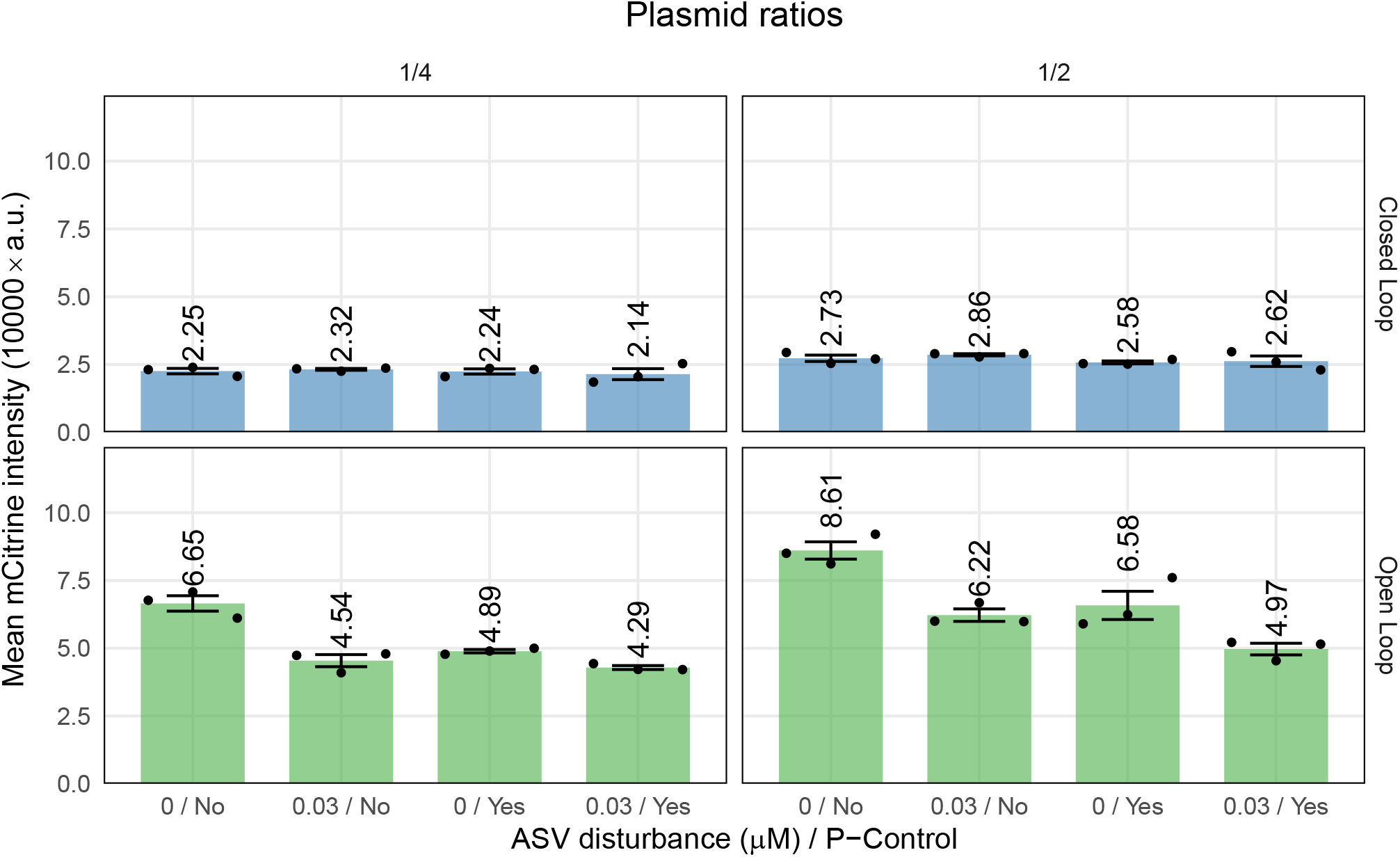
Fluorescence values shown in Figure 4(b) in arbitrary fluorescence units. The P and PI circuits were implemented by adding a negative feedback loop from L7Ae to the open- and closed-loop circuits, as shown in Figure 4(a). All circuits were perturbed by adding 30 nM of ASV. This was done for two setpoints 1/4 and 1/2 (setpoint:=activator/antisense). The data was collected 48 hours after transfection and is plotted as mean fluorescence intensity ± standard error for N = 3 replicates. The data is provided in a separate file.

**Figure A.4:**
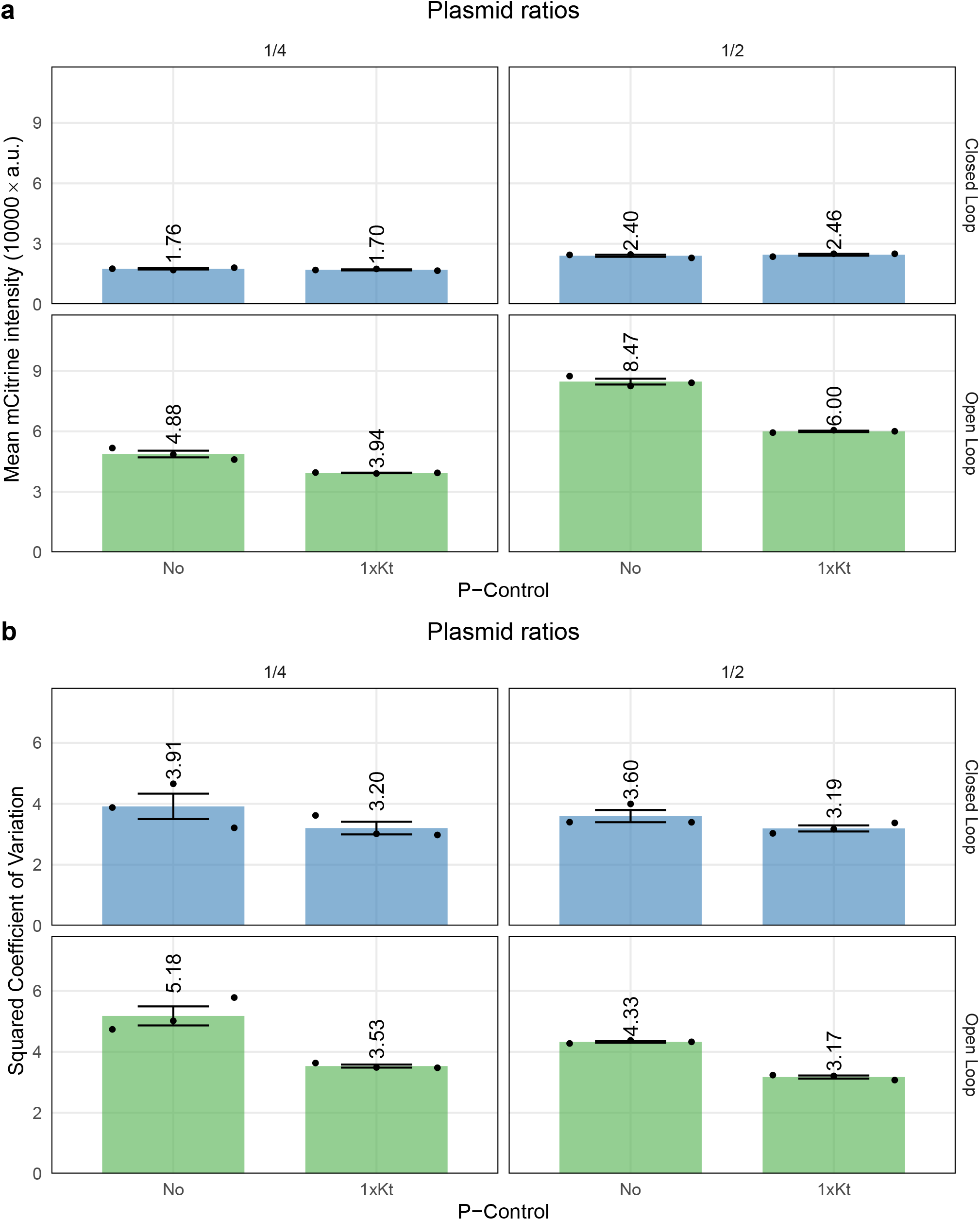
Fluorescence values shown in Figure 4(c) in arbitrary fluorescence units. This experiment was performed in a 24-well plate rather than a 96-well plate because a large sample size is required to estimate the steady-state variance accurately. The P and PI circuits were implemented by adding a negative feedback loop from L7Ae to the open- and closed-loop circuits, as shown in Figure 4(a). This was done for two setpoints 1/4 and 1/2 (setpoint:=activator/antisense). (a) Expression levels of tTA-mCitrine-SMASh are plotted as mean fluorescence intensity ± standard error for N = 3 replicates. (b) The coefficient of variation squared is shown as the mean ± standard error for N = 3 replicates per condition. The data is provided in a separate file.

**Figure A.5:**
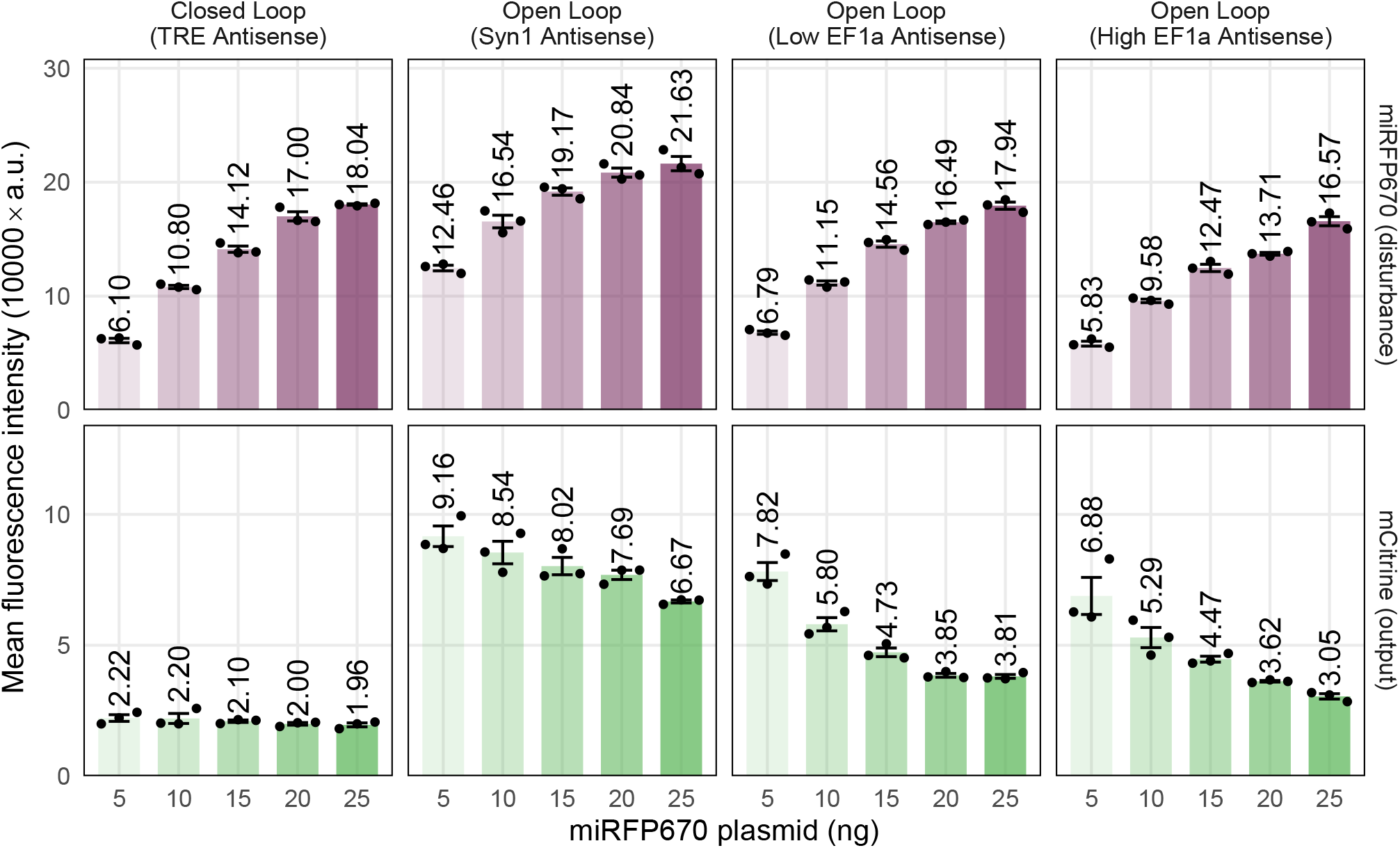
Fluorescence values of different implementations of the open-loop circuit. Besides the open- and closed-loop circuit shown in Figure 2a, another open-loop implementation, in which the antisense RNA is expressed by a strong constitutive EF1α promoter, is shown. The amount of activator plasmid was fixed among different circuits, and therefore, different levels of dsRNA formation were achieved by different amounts of antisense plasmid. All of the conditions were co-transfected with different amounts of an additional disturbance plasmid that constitutively expresses the fluorescent protein miRFP670. Expression of miRFP670 not only reflects the potential effect of dsRNA formation on gene expression but also introduces a disturbance to the amount of available resources which indirectly affects the expression levels of tTA-mCitrine-SMASh. The data was collected 48 hours after transfection and is plotted as mean fluorescence intensity ± standard error for N = 3 replicates. The data is provided in a separate file.

**Figure A.6:**
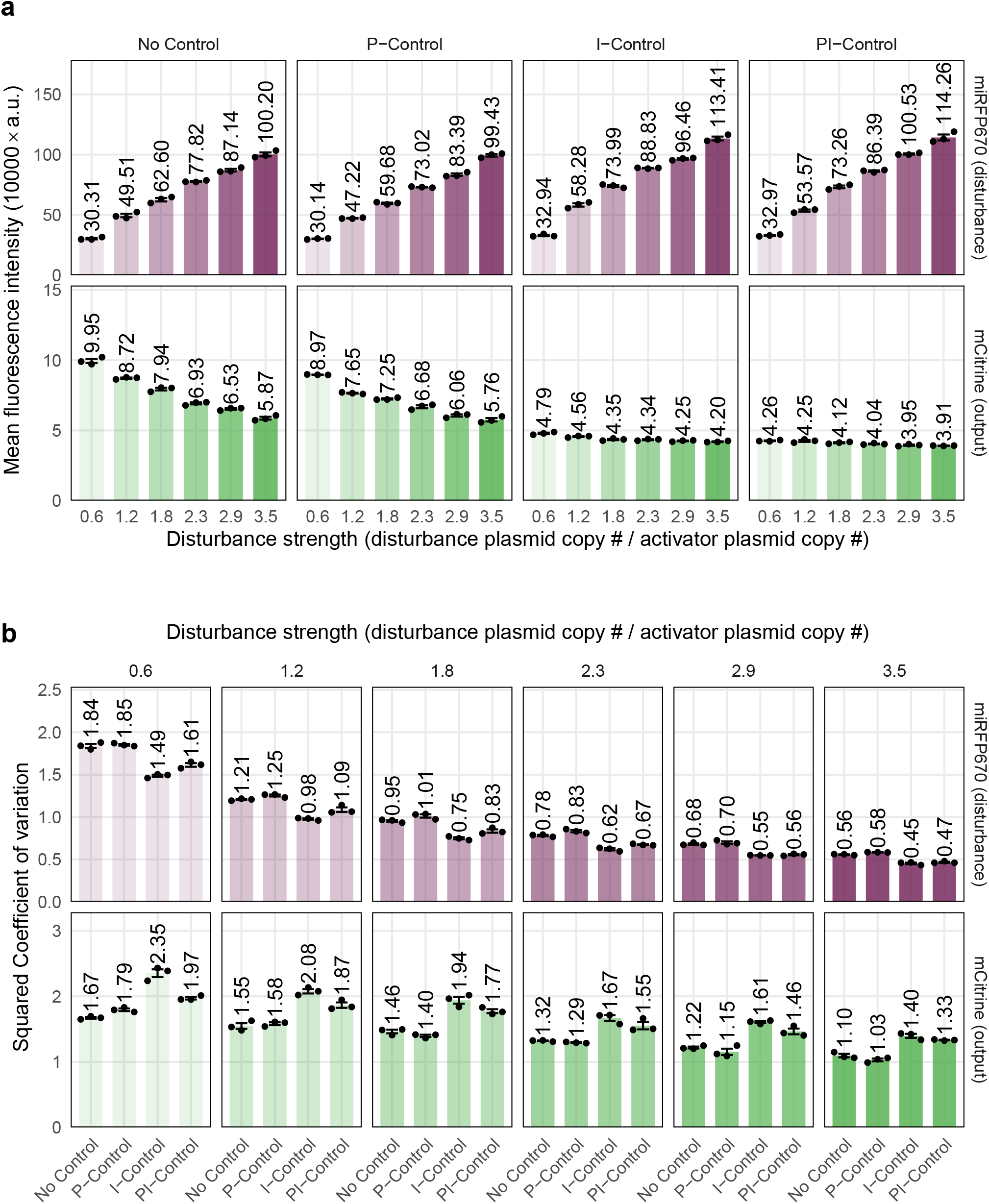
Fluorescence values shown in Figure 6 in arbitrary fluorescence units. This experiment was performed in the 24-well plate rather than 96-well plate because a large sample size is required to estimate the steady-state variance accurately. The open-loop (No Control), proportional feedback (P-Control), antithetic integral feedback (I-Control) and proportional-integral feedback (PI-Control) circuits were perturbed by co-transfecting different amounts of an additional disturbance plasmid that constitutively expresses the fluorescent protein miRFP670, as shown in Figure 6(a). The increase of miRFP670 expression introduces a disturbance to the amount of available resources which indirectly affects the expression levels of tTA-mCitrine-SMASh. The activator plasmid and antisense plasmid for all controllers were transiently transfected at a setpoint ratio of 1/2 together with disturbance strengths varying from 0.6 to 3.5. The data was collected 48 hours after transfection. (a) Expression levels of tTA-mCitrine-SMASh and miRFP670 are plotted as mean fluorescence intensity ± standard error for N = 3 replicates. (b) The coefficient of variation squared is shown as the mean ± standard error for N= 3 replicates per condition. The data is provided in a separate file.

## B Supplementary Tables

**Table B.1:** Transfection table regarding data shown in Figure 2.

|  |  |  |  |
| --- | --- | --- | --- |
| Open Loop (OL) |  |  |  |
| 100 ng total | pCH127 | pCH124 | pGLM171 |
| Plasmid ratio = 2 | 31.08 ng | 14.26 ng | 54.66 ng |
| Plasmid ratio = 1 | 15.54 ng | 14.26 ng | 70.20 ng |
| Plasmid ratio = 1/2 | 7.77 ng | 14.26 ng | 77.97 ng |
| Plasmid ratio = 1/4 | 3.89 ng | 14.26 ng | 81.86 ng |
| Plasmid ratio = 1/8 | 1.94 ng | 14.26 ng | 83.80 ng |
| Plasmid ratio = 1/16 | 0.97 ng | 14.26 ng | 84.77 ng |
| Optimen | to 10 uL |  |  |
| PEI | 0.3 uL |  |  |
| HEK293T | 12000 |  |  |
| Closed Loop (CL) |  |  |  |
| 100 ng total | pCH127 | pCH122 | pGLM171 |
| Plasmid ratio = 2 | 31.08 ng | 14.84 ng | 54.08 ng |
| Plasmid ratio = 1 | 15.54 ng | 14.84 ng | 69.62 ng |
| Plasmid ratio = 1/2 | 7.77 ng | 14.84 ng | 77.39 ng |
| Plasmid ratio = 1/4 | 3.89 ng | 14.84 ng | 81.28 ng |
| Plasmid ratio = 1/8 | 1.94 ng | 14.84 ng | 83.22 ng |
| Plasmid ratio = 1/16 | 0.97 ng | 14.84 ng | 84.19 ng |
| Optimen | to 10 uL |  |  |
| PEI | 0.3 uL |  |  |
| HEK293T | 12000 |  |  |

**Table B.2:** Transfection table regarding data shown in Figure 3. Open Loop (OL), Closed Loop (CL), With/Without Network Perturbation (NP).

|  |  |  |  |  |  |  |
| --- | --- | --- | --- | --- | --- | --- |
| Plasmid ratio = 1/2 |  |  |  |  |  |  |
| 100 ng total | pCH310 | pCH124 | pCH122 | pCH135 | pCH323 | pGLM171 |
| OL w/o NP | 7.80 ng | 14.26 ng | 0 ng | 0 ng | 2.26 ng | 75.68 ng |
| CL w/o NP | 7.80 ng | 0 ng | 14.84 ng | 0 ng | 2.26 ng | 75.1 ng |
| OL w/ NP | 7.80 ng | 14.26 ng | 0 ng | 2.41 ng | 0 ng | 75.53 ng |
| CL w/ NP | 7.80 ng | 0 ng | 14.84 ng | 2.41 ng | 0 ng | 74.95 ng |
| Optimen | to 10 uL |  |  |  |  |  |
| PEI | 0.3 uL |  |  |  |  |  |
| HEK293T | 12000 |  |  |  |  |  |
| Plasmid ratio = 1/4 |  |  |  |  |  |  |
| 100 ng total | pCH310 | pCH124 | pCH122 | pCH135 | pCH323 | pGLM171 |
| OL w/o NP | 3.90 ng | 14.26 ng | 0 ng | 0 ng | 2.26 ng | 79.58 ng |
| CL w/o NP | 3.90 ng | 0 ng | 14.84 ng | 0 ng | 2.26 ng | 79.43 ng |
| OL w/ NP | 3.90 ng | 14.26 ng | 0 ng | 2.41 ng | 0 ng | 79.00 ng |
| CL w/ NP | 3.90 ng | 0 ng | 14.84 ng | 2.41 ng | 0 ng | 78.85 ng |
| Optimen | to 10 uL |  |  |  |  |  |
| PEI | 0.3 uL |  |  |  |  |  |
| HEK293T | 12000 |  |  |  |  |  |

**Table B.3:** Transfection table regarding data shown in Figure 4(b). Open Loop (OL), Closed Loop (CL), With/Without Proportional Control (PC).

|  |  |  |  |  |  |
| --- | --- | --- | --- | --- | --- |
| Plasmid ratio = 1/2 |  |  |  |  |  |
| 100 ng total | pCH184 | pCH183 | pCH192 | pCH191 | pGLM171 |
| OL w/o NP | 8.24 ng | 0 ng | 15.26 ng | 0 ng | 76.49 ng |
| CL w/o NP | 8.24 ng | 0 ng | 0 ng | 15.84 ng | 75.91 ng |
| OL w/ NP | 0 ng | 8.24 ng | 15.26 ng | 0 ng | 76.49 ng |
| CL w/ NP | 0 ng | 8.24 ng | 0 ng | 15.84 ng | 75.91 ng |
| Optimen | to 10 uL |  |  |  |  |
| PEI | 0.3 uL |  |  |  |  |
| HEK293T | 12000 |  |  |  |  |
| Plasmid ratio = 1/4 |  |  |  |  |  |
| 100 ng total | pCH184 | pCH183 | pCH192 | pCH191 | pGLM171 |
| OL w/o NP | 4.12 ng | 0 ng | 15.26 ng | 0 ng | 80.62 ng |
| CL w/o NP | 4.12 ng | 0 ng | 0 ng | 15.84 ng | 80.04 ng |
| OL w/ NP | 0 ng | 4.12 ng | 15.26 ng | 0 ng | 80.62 ng |
| CL w/ NP | 0 ng | 4.12 ng | 0 ng | 15.84 ng | 80.04 ng |
| Optimen | to 10 uL |  |  |  |  |
| PEI | 0.3 uL |  |  |  |  |
| HEK293T | 12000 |  |  |  |  |

**Table B.4:** Transfection table regarding data shown in Figure 4(c). Open Loop (OL), Closed Loop (CL), With/Without Proportional Control (PC).

|  |  |  |  |  |  |
| --- | --- | --- | --- | --- | --- |
| Plasmid ratio = 1/2 |  |  |  |  |  |
| 500 ng total | pCH184 | pCH183 | pCH192 | pCH191 | pGLM171 |
| OL w/o NP | 41.21 ng | 0 ng | 76.32 ng | 0 ng | 382.47 ng |
| CL w/o NP | 41.21 ng | 0 ng | 0 ng | 79.22 ng | 379.57 ng |
| OL w/ NP | 0 ng | 41.21 ng | 76.32 ng | 0 ng | 382.47 ng |
| CL w/ NP | 0 ng | 41.21 ng | 0 ng | 79.22 ng | 379.57 ng |
| Optimen | to 50 uL |  |  |  |  |
| PEI | 1.5 uL |  |  |  |  |
| HEK293T | 75000 |  |  |  |  |
| Plasmid ratio = 1/4 |  |  |  |  |  |
| 100 ng total | pCH184 | pCH183 | pCH192 | pCH191 | pGLM171 |
| OL w/o NP | 20.61 ng | 0 ng | 76.32 ng | 0 ng | 403.08 ng |
| CL w/o NP | 20.61 ng | 0 ng | 0 ng | 79.22 ng | 400.18 ng |
| OL w/ NP | 0 ng | 20.61 ng | 76.32 ng | 0 ng | 403.08 ng |
| CL w/ NP | 0 ng | 20.61 ng | 0 ng | 79.22 ng | 400.18 ng |
| Optimen | to 50 uL |  |  |  |  |
| PEI | 1.5 uL |  |  |  |  |
| HEK293T | 75000 |  |  |  |  |

**Table B.5:** Transfection table regarding data shown in Figure A.5.

|  |  |  |  |  |
| --- | --- | --- | --- | --- |
| Closed Loop (TRE Antisense) |  |  |  |  |
| 100 ng total | pCH127 | pCH122 | pTTF138 | pGLM171 |
| DS = 1 | 7.77 ng | 14.84 ng | 10 ng | 67.39 ng |
| DS = 2 | 7.77 ng | 14.84 ng | 20 ng | 57.39 ng |
| DS = 3 | 7.77 ng | 14.84 ng | 30 ng | 47.39 ng |
| DS = 4 | 7.77 ng | 14.84 ng | 40 ng | 37.39 ng |
| DS = 5 | 7.77 ng | 14.84 ng | 50 ng | 27.39 ng |
| Optimen | to 10 uL |  |  |  |
| PEI | 0.3 uL |  |  |  |
| HEK293T | 12000 |  |  |  |
| Open Loop (Syn1 Antisense) |  |  |  |  |
| 100 ng total | pCH127 | pCH124 | pTTF138 | pGLM171 |
| DS = 1 | 7.77 ng | 14.26 ng | 10 ng | 67.97 ng |
| DS = 2 | 7.77 ng | 14.26 ng | 20 ng | 57.97 ng |
| DS = 3 | 7.77 ng | 14.26 ng | 30 ng | 47.97 ng |
| DS = 4 | 7.77 ng | 14.26 ng | 40 ng | 37.97 ng |
| DS = 5 | 7.77 ng | 14.26 ng | 50 ng | 27.97 ng |
| Optimen | to 10 uL |  |  |  |
| PEI | 0.3 uL |  |  |  |
| HEK293T | 12000 |  |  |  |
| Open Loop (Low EF1α Antisense) |  |  |  |  |
| 100 ng total | pCH127 | pCH218 | pTTF138 | pGLM171 |
| DS = 1 | 7.77 ng | 1.94 ng | 10 ng | 80.29 ng |
| DS = 2 | 7.77 ng | 1.94 ng | 20 ng | 70.29 ng |
| DS = 3 | 7.77 ng | 1.94 ng | 30 ng | 60.29 ng |
| DS = 4 | 7.77 ng | 1.94 ng | 40 ng | 50.29 ng |
| DS = 5 | 7.77 ng | 1.94 ng | 50 ng | 40.29 ng |
| Optimen | to 10 uL |  |  |  |
| PEI | 0.3 uL |  |  |  |
| HEK293T | 12000 |  |  |  |
| Open Loop (High EF1α Antisense) |  |  |  |  |
| 100 ng total | pCH127 | pCH218 | pTTF138 | pGLM171 |
| DS = 1 | 7.77 ng | 3.88 ng | 10 ng | 78.35 ng |
| DS = 2 | 7.77 ng | 3.88 ng | 20 ng | 68.35 ng |
| DS = 3 | 7.77 ng | 3.88 ng | 30 ng | 58.35 ng |
| DS = 4 | 7.77 ng | 3.88 ng | 40 ng | 48.35 ng |
| DS = 5 | 7.77 ng | 3.88 ng | 50 ng | 38.35 ng |
| Optimen | to 10 uL |  |  |  |
| PEI | 0.3 uL |  |  |  |
| HEK293T | 12000 |  |  |  |

**Table B.6:**
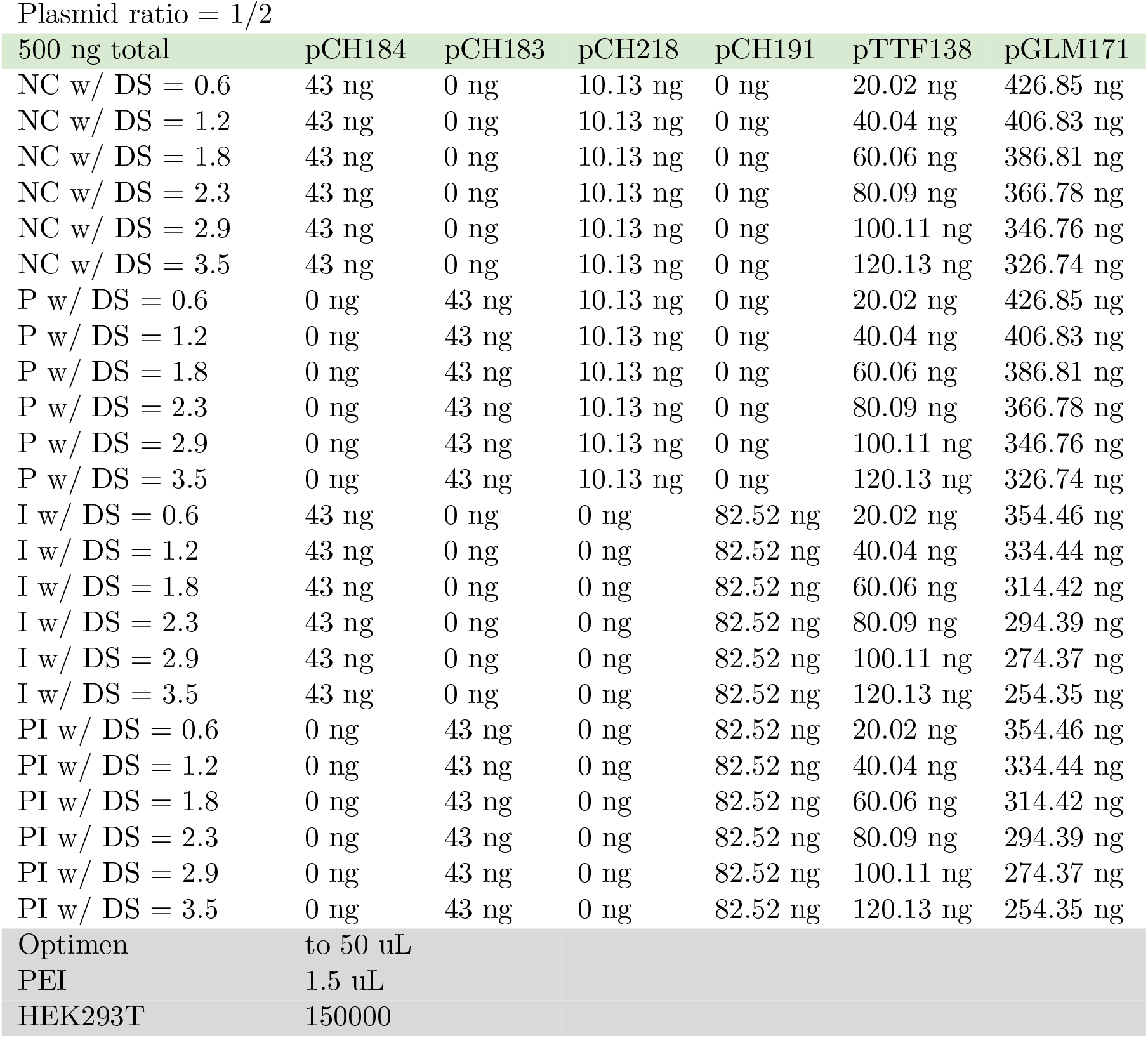
Transfection table regarding data shown in Figure 6. No Control (NC), P-Control (P), I-Control (I), PI-Control (PI), Disturbance Strength (DS).

**Table B.7:**
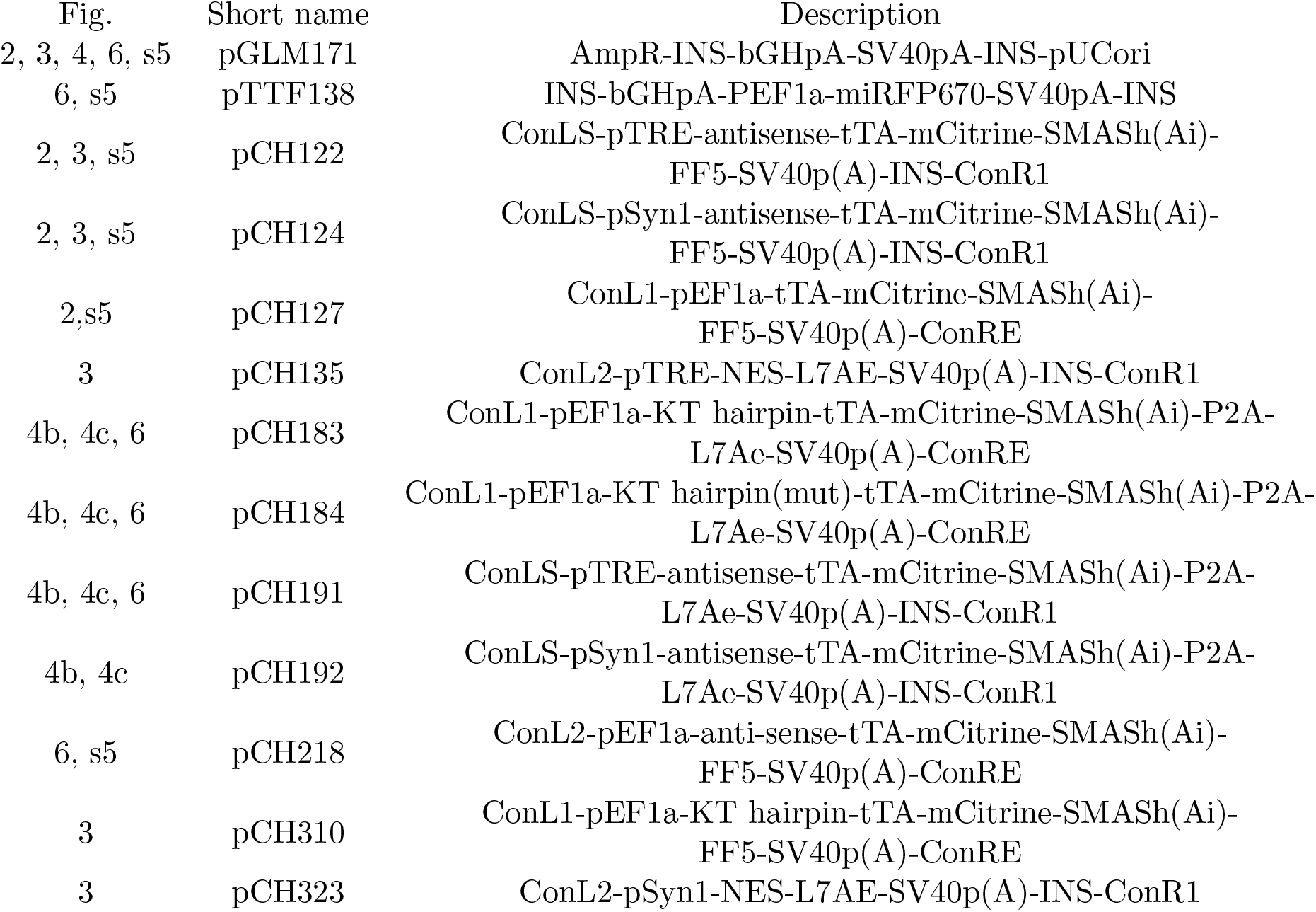
List of the plasmids used in this study. Plasmid sequences enclosed in separate file.

| Fig. | Short name | Description |
| --- | --- | --- |
| 2, 3, 4, 6, s5 | pGLM171 | AmpR-INS-bGHpA-SV40pA-INS-pUCori |
| 6, s5 | pTTF138 | INS-bGHpA-PEF1a-miRFP670-SV40pA-INS |
| 2, 3, s5 | pCH122 | ConLS-pTRE-antisense-tTA-mCitrine-SMASH(Ai)-<br>FF5-SV40p(A)-INS-ConR1 |
| 2, 3, s5 | pCH124 | ConLS-pSyn1-antisense-tTA-mCitrine-SMASH(Ai)-<br>FF5-SV40p(A)-INS-ConR1 |
| 2,s5 | pCH127 | ConL1-pEF1a-tTA-mCitrine-SMASH(Ai)-<br>FF5-SV40p(A)-ConRE |
| 3 | pCH135 | ConL2-pTRE-NES-L7AE-SV40p(A)-INS-ConR1 |
| 4b, 4c, 6 | pCH183 | ConL1-pEF1a-KT hairpin-tTA-mCitrine-SMASH(Ai)-P2A-L7Ae-SV40p(A)-ConRE |
| 4b, 4c, 6 | pCH184 | ConL1-pEF1a-KT hairpin(mut)-tTA-mCitrine-SMASH(Ai)-P2A-L7Ae-SV40p(A)-ConRE |
| 4b, 4c, 6 | pCH191 | ConLS-pTRE-antisense-tTA-mCitrine-SMASH(Ai)-P2A-L7Ae-SV40p(A)-INS-ConR1 |
| 4b, 4c | pCH192 | ConLS-pSyn1-antisense-tTA-mCitrine-SMASH(Ai)-P2A-L7Ae-SV40p(A)-INS-ConR1 |
| 6, s5 | pCH218 | ConL2-pEF1a-anti-sense-tTA-mCitrine-SMASH(Ai)-<br>FF5-SV40p(A)-ConRE |
| 3 | pCH310 | ConL1-pEF1a-KT hairpin-tTA-mCitrine-SMASH(Ai)-<br>FF5-SV40p(A)-ConRE |
| 3 | pCH323 | ConL2-pSyn1-NES-L7AE-SV40p(A)-INS-ConR1 |

## C Mathematical Modeling of the Circuit in Figure 2

Consider the circuit depicted in Figure 2(a) where an RNA-based molecular realization of the antithetic integral controller regulates the production of a particular protein of interest, namely the transcription factor tTA-mCitrine. The circuit can operate in either open loop or closed loop. We first present a detailed (mechanistic) mathematical model and then carry out a model reduction technique that allows us to analyze the steady-state behavior of the output protein. Finally, we provide the technical details of properly calibrating the model to the experimental data.

### C.1 Full Model Description

A detailed biochemical reaction network that describes the interactions between the various biochemical species, depicted in Table C.8, is given in Table C.9. These tables are sufficient to provide a mathematical model for the circuit in Figure 2(a). The open-loop circuit can be obtained by setting *a*_2_ = 0 (in Table C.9) thus preventing the transcriptional activator **A** from binding to the promoter of the anti-sense gene 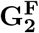. Note that if **A** and **B** are two species, then **A:B** is understood to be the complex formed when **A** and **B** are bound together.

**Table C.8:** List of Biochemical Species

| Species | Symbol | Species | Symbol |
| --- | --- | --- | --- |
| Activator Gene | $\mathbf{G}_1$ | Antisense Gene (Free) | $\mathbf{G}_2^F$ |
| Sense mRNA | $\mathbf{Z}_1$ | Anti-sense RNA | $\mathbf{Z}_2$ |
| Asunaprevir Disturbance (Free) | $\mathbf{D}^F$ | tTA:mCitrine:SMASh | $\mathbf{X}_1$ |
| $\mathbf{X}_1:\mathbf{D}^F$ | $\mathbf{X}_1^*$ | tTA:mCitrine | $\mathbf{X}_2$ |
| Dimerized tTA:mCitrine | $\mathbf{A}$ | $\mathbf{G}_2^F:\mathbf{A}$ | $\mathbf{G}_2^*$ |

**Table C.9:**
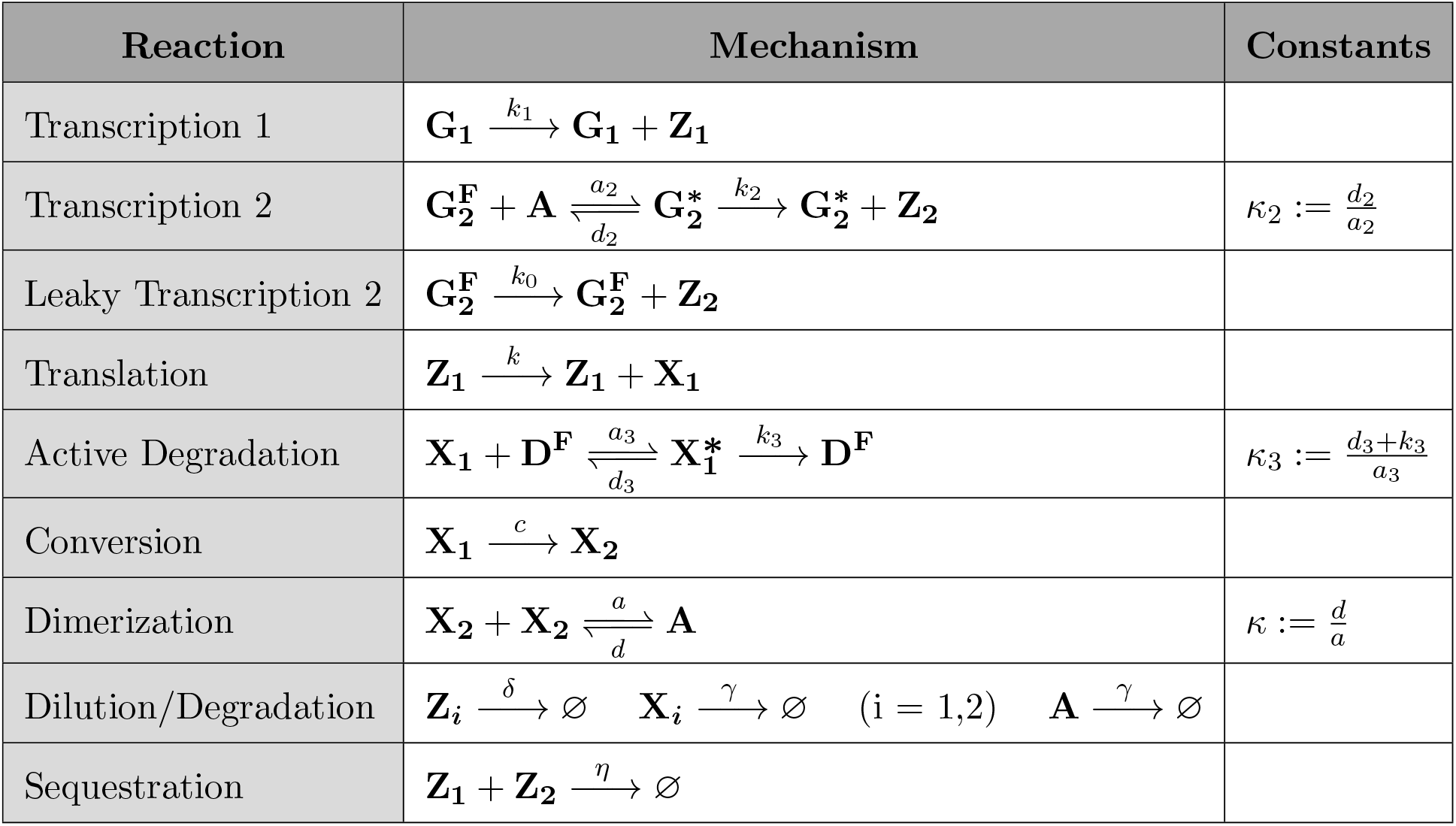
List of Biochemical Reactions

### C.2 Model Reduction

In this section, the full model given in Table C.9 is mathematically reduced to the model described schematically in Figure C.7(a) and mathematically in Figure C.7(b).

**Figure C.7:**
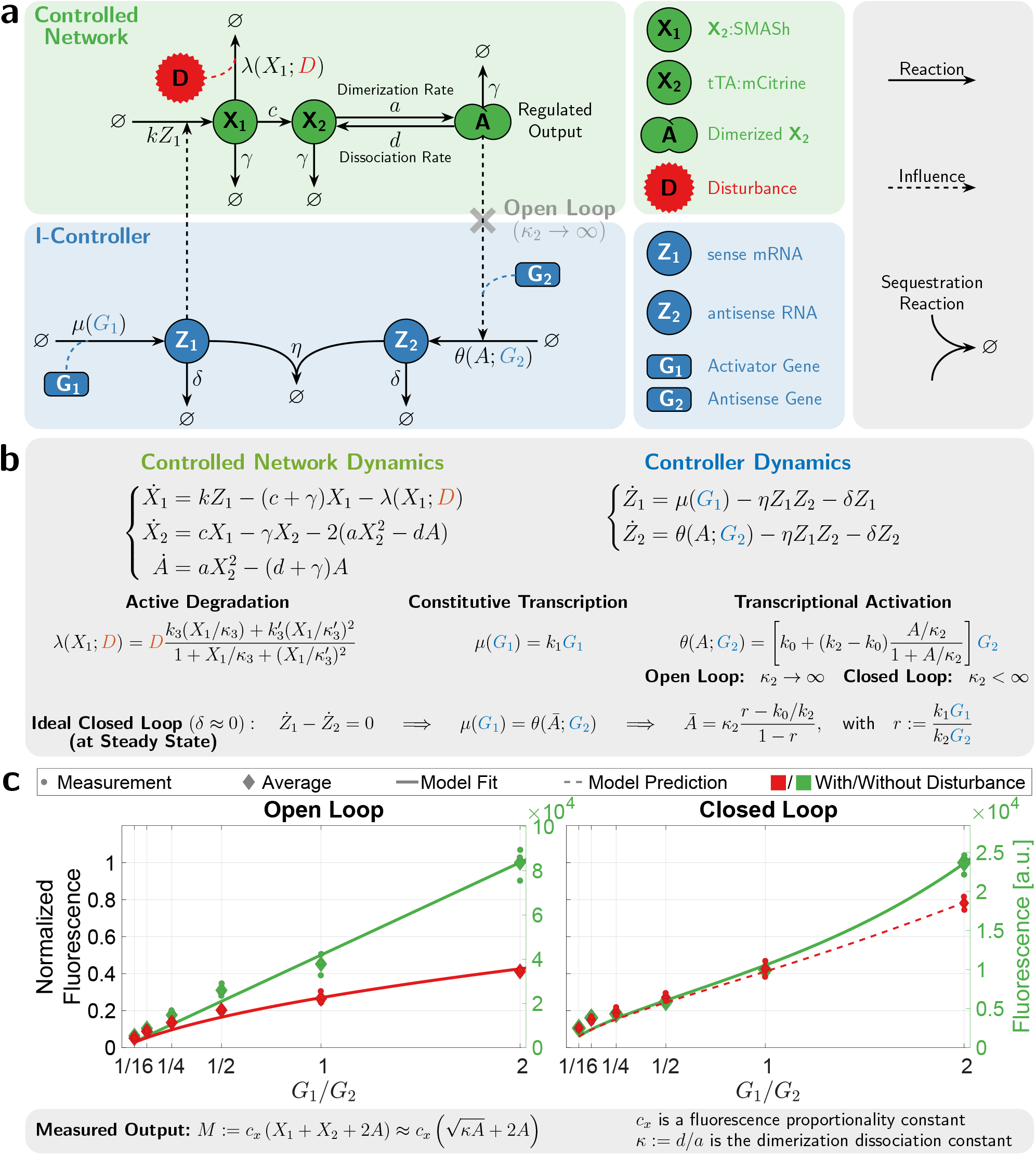
Mathematical Modeling of the I-Circuit in Figure 2. **(a)/(b) Schematic/Mathematical Description of the Reduced Model.** This is a special case of the compact model presented in Figure 5(a) where the proportional controller and network perturbation are removed to model only the integral control action. The model for the open-loop circuit is obtained by setting *κ*_2_ → ∞, where *κ*_2_ is the dissociation constant of **A** from **G_2_**. This removes the the feedback from the regulated output **A** since *θ*(*A*; **G**_2_) becomes *k*_0_*G*_2_. In the ideal operation of the antithetic integral controller, where the dilution rate *δ* is negligible with respect to the other rates of the controller, the regulated output **A** has a steady-state concentration, denoted by *Ā*, that is independent of the controlled network parameters. This ensures robust perfect adaptation of the regulated output to external disturbances such as **D**. **(c)Model Calibration to Experimental Data.** This panel is the same as Figure 5(b), but the x-axis is plotted on a linear scale here to examine the concavity of the curves.

Note that Figure C.7 is a special case of Figure 5(a) in the main text, where there is only integral control and no network perturbation. The model reduction procedure is based on the following assumption:

#### Assumption 1.

The binding/unbinding reactions are fast.

Assumption 1 allows us to exploit a time-scale separation principle based on the fact that the (un)binding reactions are much faster than the other reactions in the system. As a result, a Quasi-Steady-State Approximation (QSSA) is applied.

Now, we show the mathematical derivation of the reduced model. The conservation laws are given in terms of the total concentrations of bound and free antisense gene and ASV denoted by *G*_2_ and *D*, respectively. That is, we have

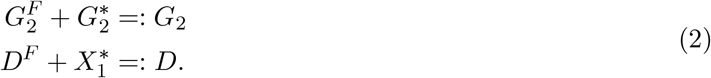

Note that *G*_1_, *G*_2_ and *D* are constants and are considered to act as external inputs and disturbance to the circuit. Since the binding reactions are much faster than the other reactions in the network (Assumption 1), one can invoke the Quasi-Steady-State Approximation (QSSA) as follows

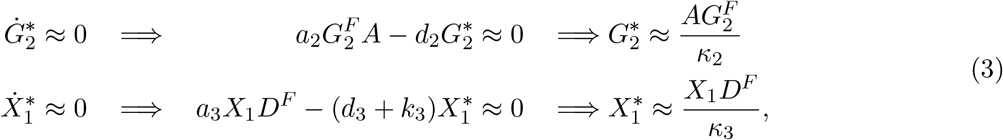

where the dissociation and Michaelis-Menten constants, **κ**_2_ and **κ**_3_, are given in Table C.9. By substituting the quasi-steady-state approximation of 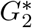 in the conservation law 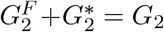, we obtain the following expressions

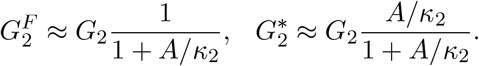

Similarly, by substituting the quasi-steady-state approximation of 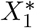 in the conservation law 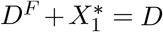, we obtain

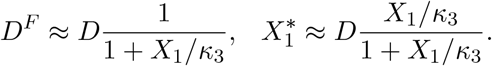

Equipped with the quasi-steady-state approximations, we can now write down a set of *Ordinary Differential Equations (ODEs)* that describe the evolution of *X*_1_, *X*_2_, *A, Z*_1_ and *Z*_2_.

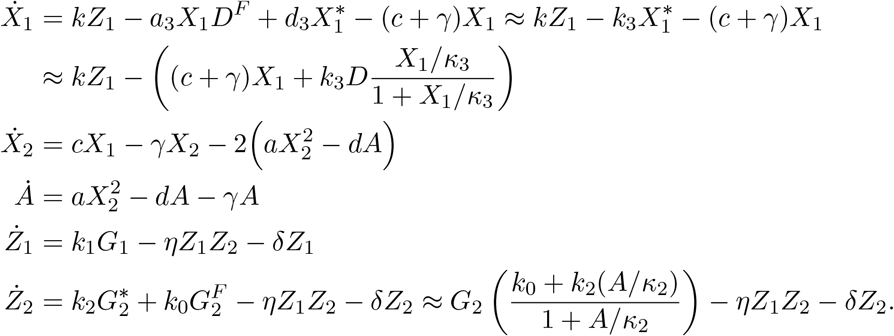

Finally, the dynamics of the reduced model can be more compactly written as

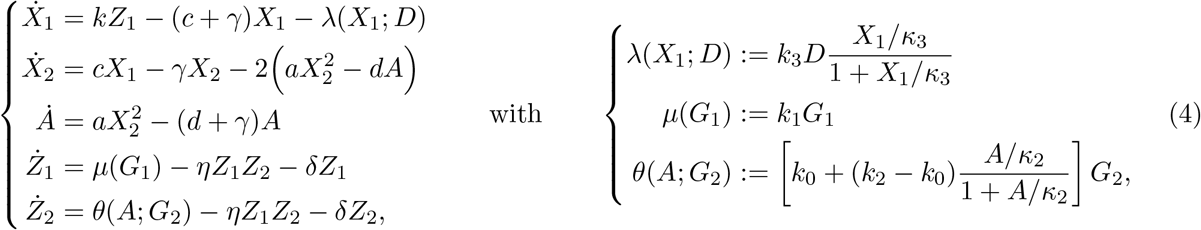

where *k*_0_ ≪ *k*_2_ since leaky transcription is usually much slower than activated transcription. Note that in the open-loop circuit, only the leaky transcription reaction can occur, that is *a*_2_ = 0. As a result, *κ*_2_ → ∞, and therefore the function *θ* becomes independent of *A*, i.e. *θ*(*A*; *G*_2_) = *k*_0_*G*_2_.

The only difference between the dynamics given in (4) and Figure C.7(b) lies in the active degradation function *λ*. In fact, by setting 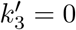 and 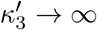 in Figure C.7(b), we obtain (4). The active degradation function of Figure C.7(b) is, in general, of higher order and involves squared terms 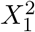. It turns out that this higher order function is necessary to fit the data properly (refer to Supplementary Information C.3 for a detailed explanation). The mechanism underlying this higher-order active degradation function is explained in the subsequent section.

### C.3 Higher Order Active Degradation

The active degradation function *λ*(*X*_1_; *D*) in (4) takes the form of a hill function multiplied by the (disturbance) Asunaprevir concentration *D*. We now consider the more general hill function of Figure C.7(b) which is given by

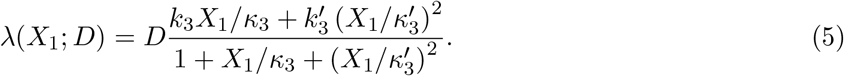

This function has a higher order (hill coefficient) since it involves squared terms 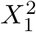. We show next how this hill function can mechanistically arise from the interactions of the various species in the circuit.

In addition to the Active Degradation reaction given in Table C.9, we allow **X_1_** to dimerize to form the complex **A_1_** (Dimerized tTA:mCitrine:SMASh) which can also be actively degraded by **D**. These additional mechanisms are modeled by appending the previous model with the additional active degradation and dimerization reactions listed in Table C.10. Note that, theoretically, **A_1_** can still release the SMASh tag at some rate *c′* and/or may still be able to initiate transcription of **G_2_** at some rate 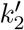. However, we assume that the bulk dimer **A_1_** is very unstable and tends to dissociate at a rate 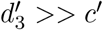 and its transcription rate is much slower than that of **A**. As a result, *c′* and 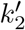 can be neglected and thus the corresponding reactions are not listed in Table C.10.

**Table C.10:** Additional Biochemical Reactions for Higher Order Active Degradation

| Reaction | Mechanism | Constants |
| --- | --- | --- |
| Dimerization 2 | $\mathbf{X}_1 + \mathbf{X}_1 \xrightleftharpoons[d'_1]{a'_1} \mathbf{A}_1$ | $\kappa'_1 := \frac{d'_1}{a'_1}$ |
| Active Degradation 2 | $\mathbf{A}_1 + \mathbf{D}^F \xrightleftharpoons[d'_3]{a'_3} \mathbf{A}_1^* \xrightarrow{k'_3} \mathbf{D}^F$ | $\kappa'_3 := \frac{d'_3 + k'_3}{a'_3}$ |

Now, we show the mathematical derivation of the higher order active degradation function *λ*(*X*_1_; *D*). The mathematical procedure is, once again, based on Assumptions 1; however, an additional assumption is added here as well.

#### Assumption 2.

The concentration of the free ASV molecules D^F^ is low.

This assumption means that the majority of the ASV molecules are in their bound state. The conservation law that can be seen from Tables C.9 and C.10 is given by

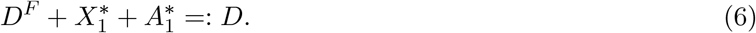

This replaces the conservation law for *D* given in (2) since the species 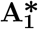 is a complex formed of **A_1_** and **D^F^**. With Assumption 1 in mind, one can invoke the Quasi-Steady-State Approximation (QSSA) as follows

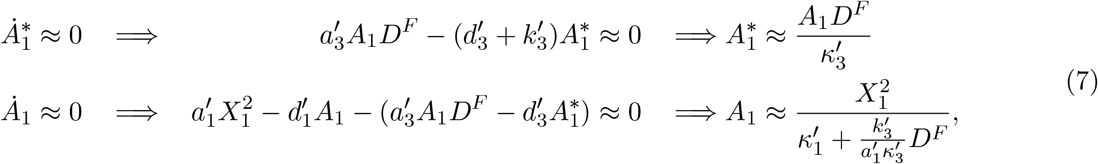

where the constants 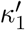 and 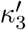 are given in Table C.10. By substituting the quasi-steady-state approximations of 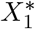 from (3) and 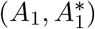 from (7) in the conservation law 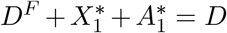, while invoking Assumption 2 (more precisely 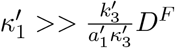), we obtain

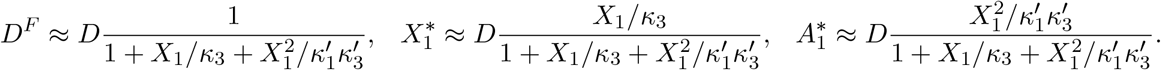

Equipped with the quasi-steady-state approximations, we can now update the *Ordinary Differential Equation (ODE)* that describes the evolution of *X*_1_.

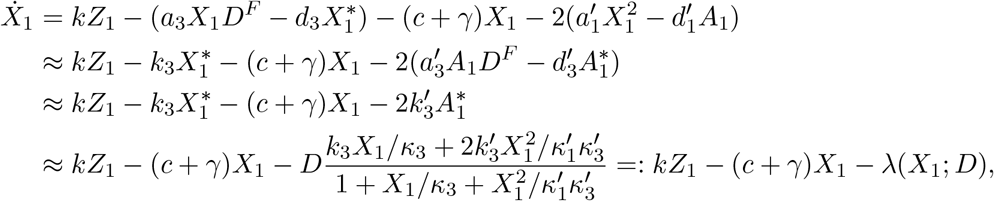

and thus the active degradation function *λ* takes the intended form given in (5) (with slight abuse of notation) and shown in Figure C.7(b).

### C.4 Mathematical Model of the Measured Output: Fluorescence

Let *M^i^* (*G*_1_, *D*) denote the measured fluorescence at a given concentration of the activator gene *G*_1_ and drug disturbance *D*, where *i* = *o* corresponds to open-loop measurements while *i* = *c* corresponds to closed-loop measurements. Note that *G*_2_ is held constant throughout the paper, and thus the explicit dependence of the measured fluorescence *M* on *G*_2_ is not emphasized here. Fluorescence is emitted from all molecules containing mCitrine, that is species **X_1_, X_2_** and **A** (see Figure C.7(a)). Hence we have

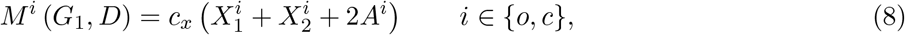

where *c_x_* is a proportionality constant that maps concentrations (in nm) to fluorescence (in a.u.). Recall that the dynamics of *X*_1_, *X*_2_ and *A* (in both open and closed loop) are given by

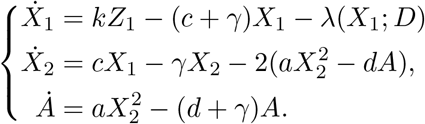

At steady state, we have 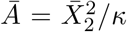 with *κ*:= (*d* + *γ*)/*a*. which implies that 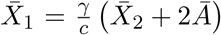. This implies that the measurement at steady state is given by

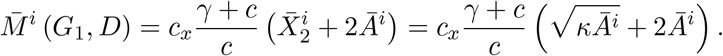

This equation links the measured fluorescence at steady state, 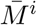, to the steady-state concentration of the regulated output *Ā^i^* for both open- and closed-loop circuits. In fact, this equation is also approximately valid transiently (not only at steady state) under the following assumption.

#### Assumption 3.

Transcription is much slower than releasing the SMASh tag.

This assumption is reasonable since **X_1_** is very unstable and tends to release the SMASh tag very quickly (i.e. *c* is large). Under this assumption, a quasi-steady-state argument can be used to obtain the following approximations 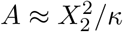 and 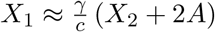 which yield the same measurement equation at any time *t* (not just at steady state). Furthermore, we can drop the factor (*γ* + *c*)/*c* since *c* ≫ *γ* to obtain the expression shown in the bottom of Figure C.7(c) and Figure 5(a).

### C.5 Model Calibration to the Experimental Data

In this section, we calibrate the model depicted in Figures C.7(a) and (b) to fit the experimentally collected data at steady state.

#### C.5.1 Mathematical Representation of the Data

The experiments that are carried out allows us to obtain the data set visualized in Figure C.8. The mathematical notation for the available data is described in Table C.11.

**Table C.11:** Data Representation: Set of inputs *G*_1_ and measured outputs 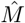 for the open- and closed-loop circuits with and without disturbance. *N°* (resp. *N^c^*) denotes the number of different concentrations of **G_1_** that are applied to the open-loop (resp. closed-loop) circuit. For each 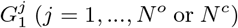, four measurements 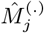 are obtained. 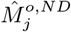 (resp. 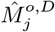) denotes the measurement for the open-loop circuit without (resp. with) disturbance; whereas, 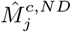 (resp. 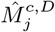) denotes the measurement for the closed-loop circuit without (resp. with) disturbance. Constant disturbances *D* = 30 nM are applied and the concentration of the antisense plasmid *G*_2_ = 0.004 pmol is kept constant throughout all the experiments. Note that the measurements 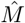 represent the average of the experimentally obtained triplicates.

| | No Disturbance ($D = 0$ nM) | With Disturbance ($D = 33$ nM) |
| --- | --- | --- |
| Open Loop | $\left\{ G_1^j, \hat{M}_j^{o,ND} \right\}_{j=1}^{N^o}$ | $\left\{ G_1^j, \hat{M}_j^{o,D} \right\}_{j=1}^{N^o}$ |
| Closed Loop | $\left\{ G_1^j, \hat{M}_j^{c,ND} \right\}_{j=1}^{N^c}$ | $\left\{ G_1^j, \hat{M}_j^{c,D} \right\}_{j=1}^{N^c}$ |

#### C.5.2 Necessity of Higher Order Active Degradation

We select the higher order active degradation function introduced in Supplementary Information C.3 because it will be shown that the negative concavity of the fitted curve in the open-loop setting with disturbance (see Figure C.7(c)) cannot be captured by a simple first order hill function *λ*.

The fixed point of the dynamics in the open-loop setting, denoted by 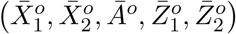 satisfies the following set of nonlinear algebraic equations.

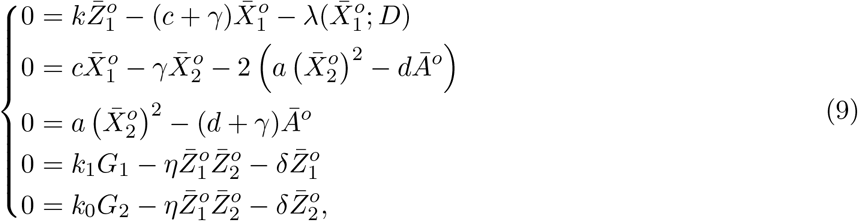

**Figure C.8:**
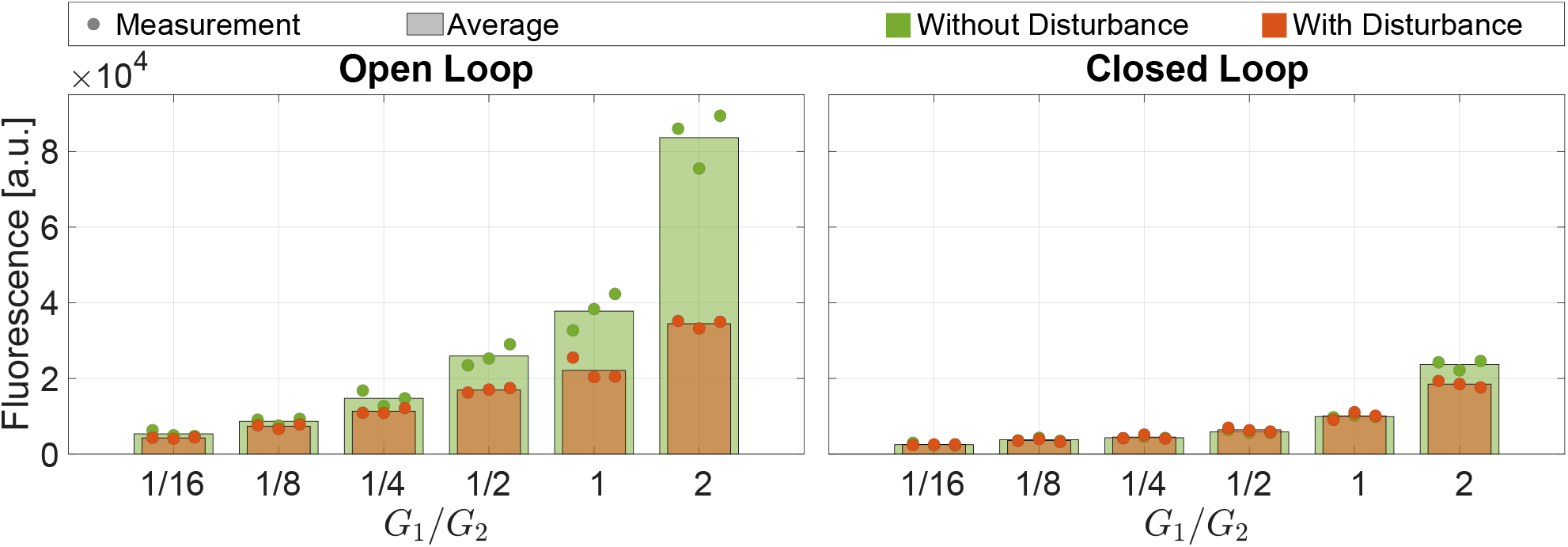
Data used for fitting the circuit realizing an antithetic integral controller. The fluorescence data are obtained for a wide range of plasmid ratios *G*_1_/*G*_2_, for open/closed loop settings, and with/without disturbance by fixing *G*_2_ = 0.004 pmol and sweeping *G*_1_ accordingly.

and the measured output denoted by 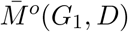 is given by

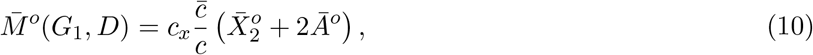

where 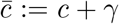. To analyze the concavity of 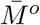 as a function of *G*_1_, we study the sign of the second derivative 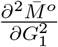. First, observe that using the second and third equations in (9), the measured output can be rewritten as

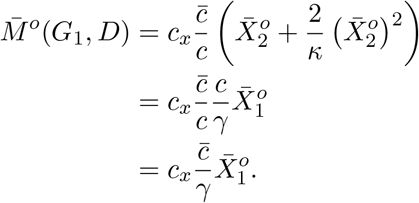

Hence the sign of 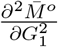 is the same as that of 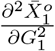 which we derive next. Taking the second derivative of the first equation in (9) with respect to *G*_1_ yields

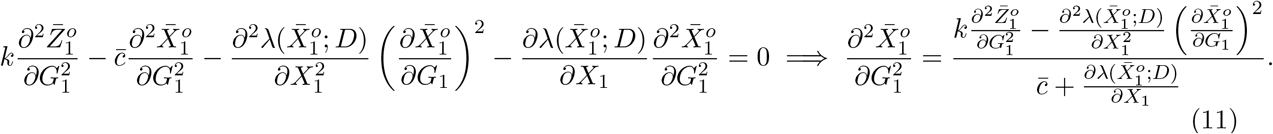

Next we derive an expression for 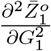. Subtracting the last two equations in (9) from each other yields

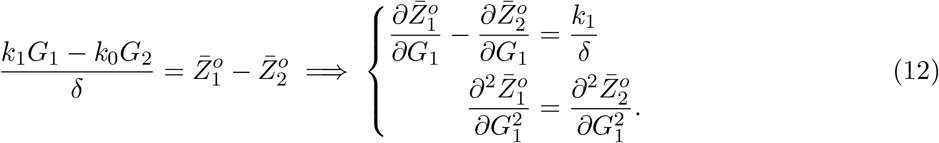

By taking the first derivative of the fourth equation in (9) with respect to *G*_1_ and exploiting (12), we obtain an expression for 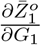 given by

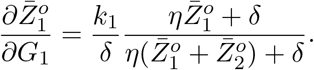

Furthermore, by taking the second derivative of the fourth equation in (9) with respect to *G*_1_ and exploiting (12), we obtain an expression for 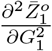 given by

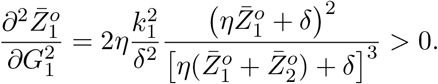

Therefore, observe using (11), that as long as λ is an increasing function of 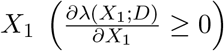 with negative concavity 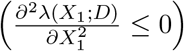, we have 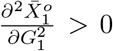. Therefore for a first order active degradation function 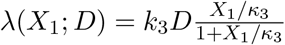, we have that 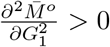 and, as a result, it cannot capture the negative concavity of the measurements in the open-loop setting with disturbance shown in Figure C.7(c). On the other hand a second order active degradation function *λ* given in (5) is capable of changing the concavity and hence is adopted in the paper.

#### C.5.3 Choice of Parameter Groups

Consider the circuit depicted in Figures C.7(a) and (b). The measured output for the open- (*i* = *o*) and closed-loop (*i* = *c*) settings is given by

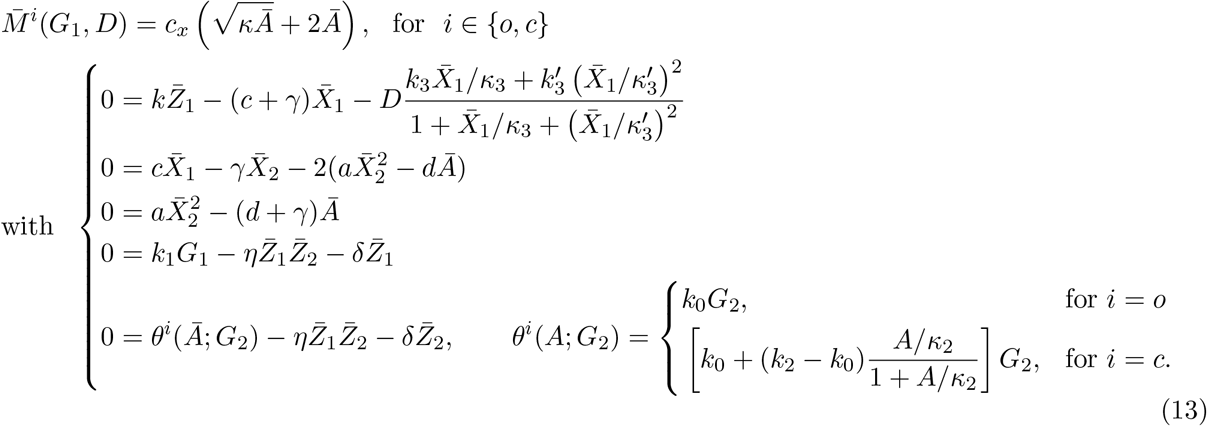

The system model has 15 parameters: 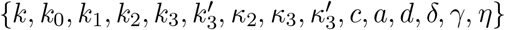 to be calibrated to the data. Furthermore, the measurement equation has an additional parameter c*x* to be calibrated as well and thus summing up to 16 total parameters. However, steady-state measurements cannot uniquely identify all of those parameters. For this reason, we carry out a suitable choice of re-parameterization to obtain a minimal number of (aggregated) parameter groups that can be uniquely identified from the steady-state measurements. Particularly, define the following parameter groups

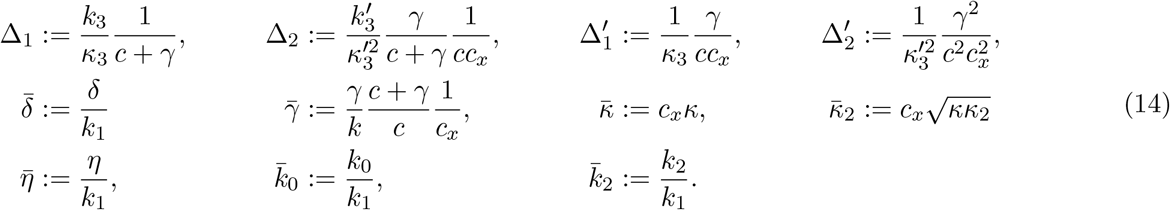

and the following transformed variables

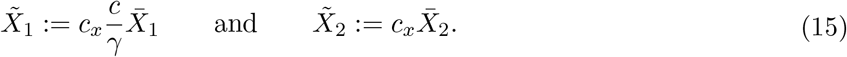

Then the steady-state measurements can be rewritten in terms of the parameter groups and transformed variables as

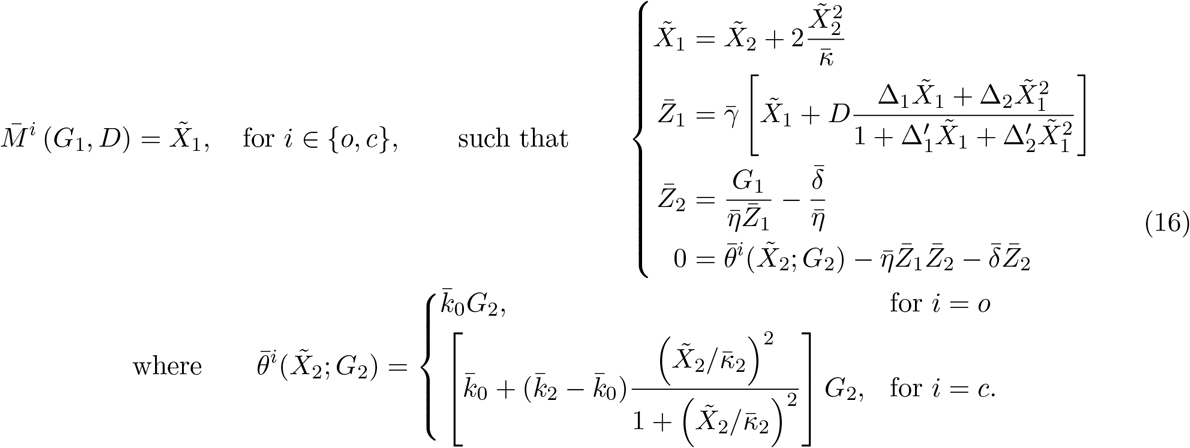

Note that *i* = *o* and *i* = *c* correspond to the open- and closed-loop settings, respectively. Hence given the input *G*_1_ and the disturbance *D*, one can use (16) to compute the mCitrine measurement in the open- and closed-loop settings. To do so, one has to solve the set of nonlinear algebraic equations. This is done in Matlab by recasting the set of nonlinear algebraic equations as a single but high order polynomial in 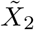 using the symbolic toolbox and then solving the polynomial using the command “roots”. This allows us to solve the system of equations more efficiently (by computing eigenvalues of a companion matrix associated with the obtained polynomial) without requiring an initial guess as in Newton-Raphson-like methods. This is particularly important since model calibration may require solving this system of equations thousands of times. We close this section by observing that we have now reduced the parameters to be calibrated down to 11 (as compared to 16).

#### C.5.4 Model Calibration Steps

The model calibration is carried out in four steps to avoid over-fitting. In the first step, the parameters of the model for the open-loop circuit in the absence of disturbance (*D* = 0) are fit to the data 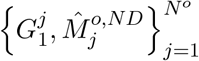 by solving the following optimization problem for 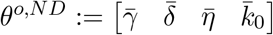.

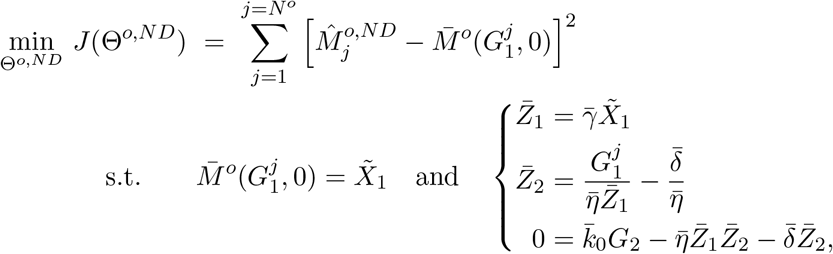

where *G*_2_ = 0.004 pmol. Note that this system of equations can be rewritten in terms of 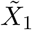 explicitly as a second degree polynomial.

In the second step, we first fix the parameters that are obtained from the previous fit. Then, the parameters of the model for the closed-loop circuit in the absence of disturbance (*D* = 0) are fit to the data 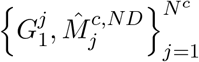 by solving the following optimization problem for 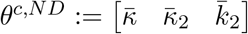.

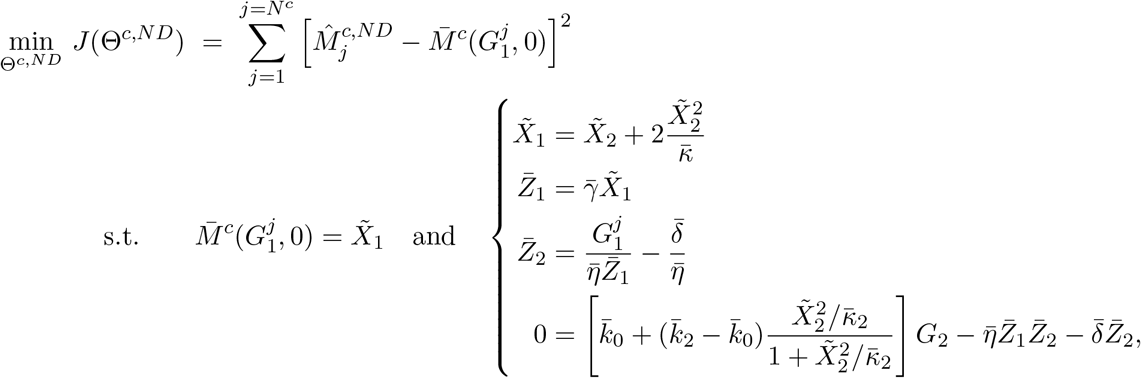

where *G*_2_ = 0.004 pmol. Note that the last equation can be rewritten in terms of 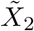 explicitly as a sixth degree polynomial.

In the third step, we first fix the parameters that are obtained from the previous fits. Then, the parameters of the model for the open-loop circuit in the presence of disturbance (*D* = 33 nm) are fit to the data 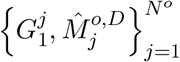 by solving the following optimization problem for 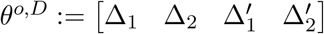.

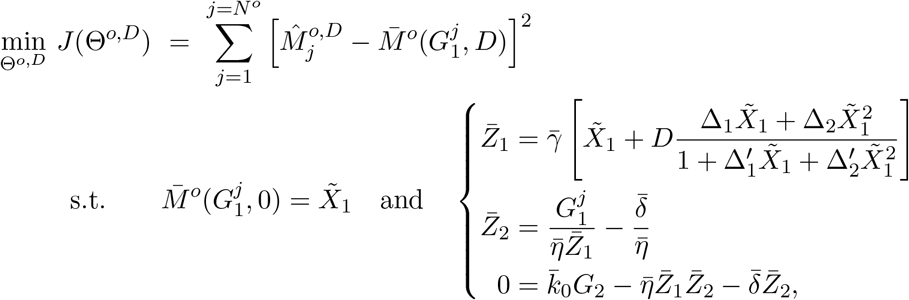

where *G*_2_ = 0.004 pmol, *D* = 33 nm. Note that the last equation can be rewritten in terms of 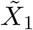 explicitly as a sixth degree polynomial.

In the last step, we already have all the parameter groups, and thus we use them to mathematically predict the measurements for the closed-loop circuit with disturbance (*D* = 33 nm). This is done by using (16) where the system of equations can be rewritten as a polynomial in 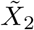 of degree 13.

##### Estimated Parameter Groups

The model fit and prediction are shown in Figures 5(b) and C.7(c), where the optimally estimated parameter groups are given by

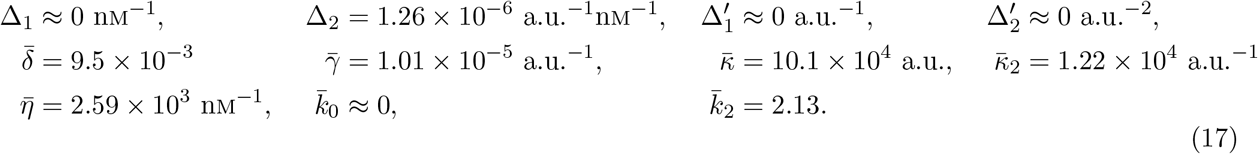

The estimated parameter groups suggest that leaky transcription of the antisense gene is negligible. They also suggest that the active degradation function is approximately purely quadratic in *X*_1_ that is 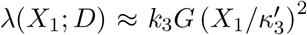 which means that the degradation of the dimerized tTA:mCitrine:SMASh dominates the degradation of the monomer.

## D Mathematical Modeling of the Circuit in Figure 3

Consider the circuit depicted in Figure 3(a) which is similar to the circuit in Figure 2(a) that is mathematically modeled and analyzed in Supplementary Information C. The difference here lies in the additional plasmid encoding a gene that expresses an RNA-binding protein capable of inhibiting the translation of the sense mRNA. We first describe the additional mechanistic interactions that are introduced by this gene, and then carry out a model reduction technique that allows us to analyze the steady-state behavior of the output protein. Finally we provide the technical details of fitting the model to the experimental data.

### D.1 Full Model Description

A detailed biochemical reaction network that models the dynamics of the circuit in Figure 3(a) can be obtained by appending Tables C.8 and C.9 (that model the circuit of Figure 2(a)) by the list of biochemical species and reactions given in Tables D.12 and D.13, respectively. These species and reactions describe the biochemical reaction sub-network that is introduced by the additional gene encoding for NES-L7Ae.

**Table D.12:**
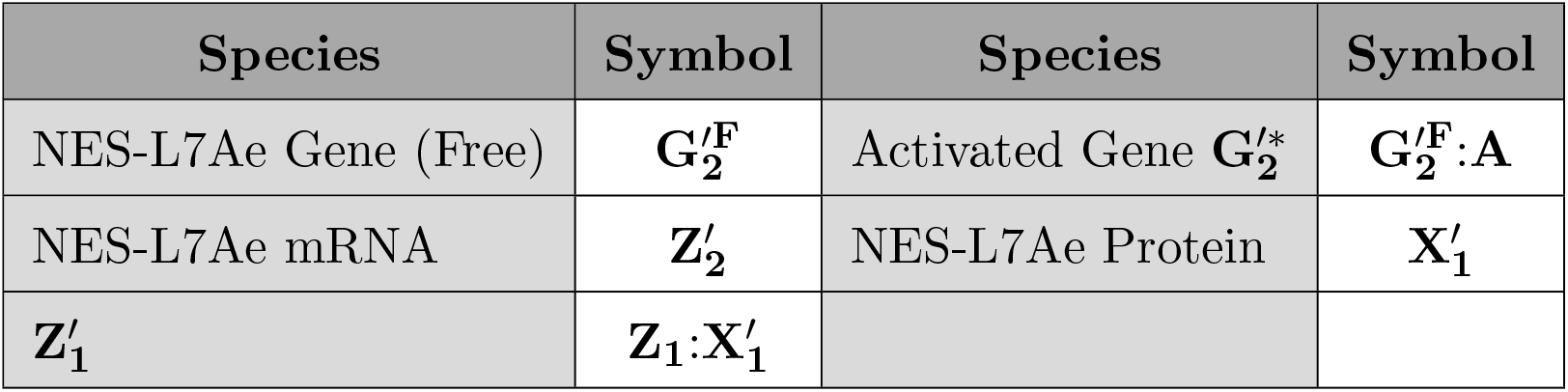
List of Additional Biochemical Species

**Table D.13:**
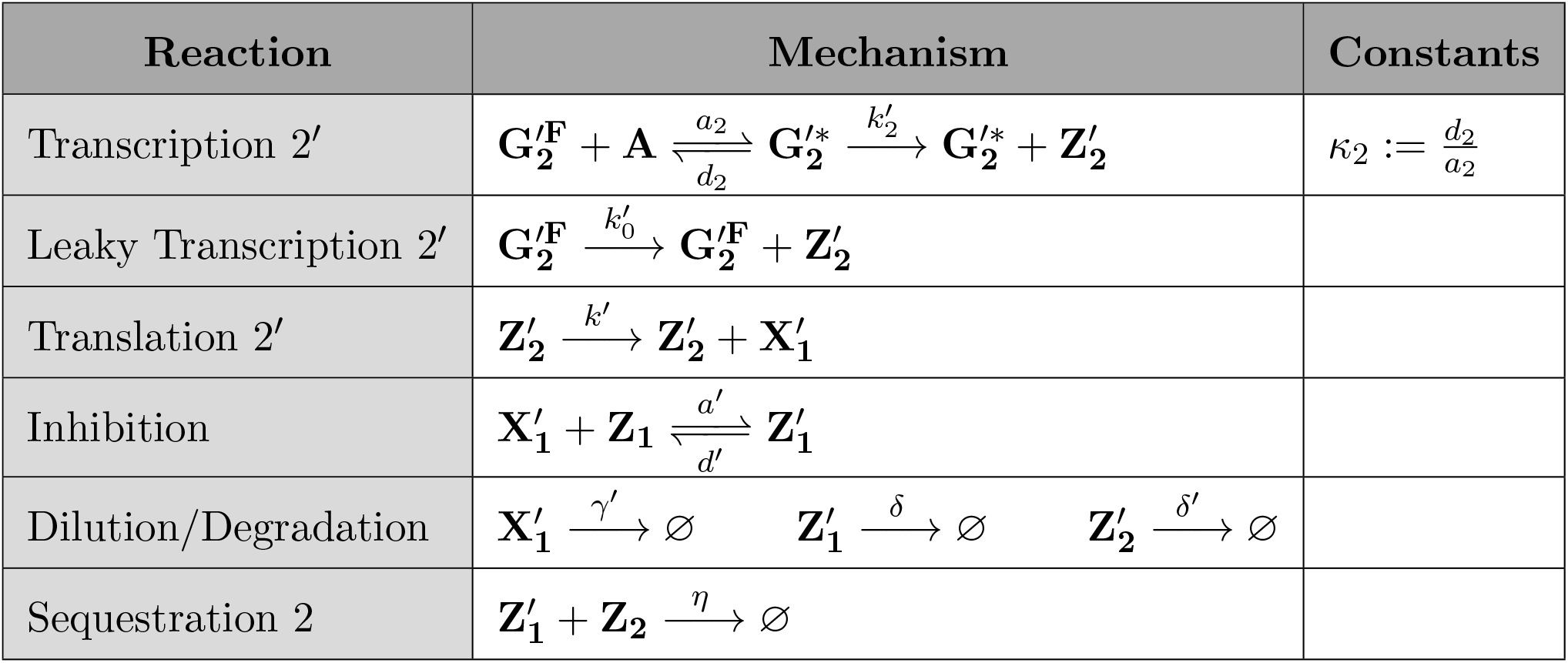
List of Additional Biochemical Reactions

### D.2 Model Reduction

In this section, the full model given in Tables C.9 and D.13 is mathematically reduced to the model described schematically in Figure D.9(a) and mathematically in Figure D.9(b). Note that Figure D.9 is a special case of Figure 5(a) in the main text, where there is only integral control but with network perturbation. The model reduction procedure is based on the reduced model that is obtained previously in Supplementary Information C.2.

**Figure D.9:**
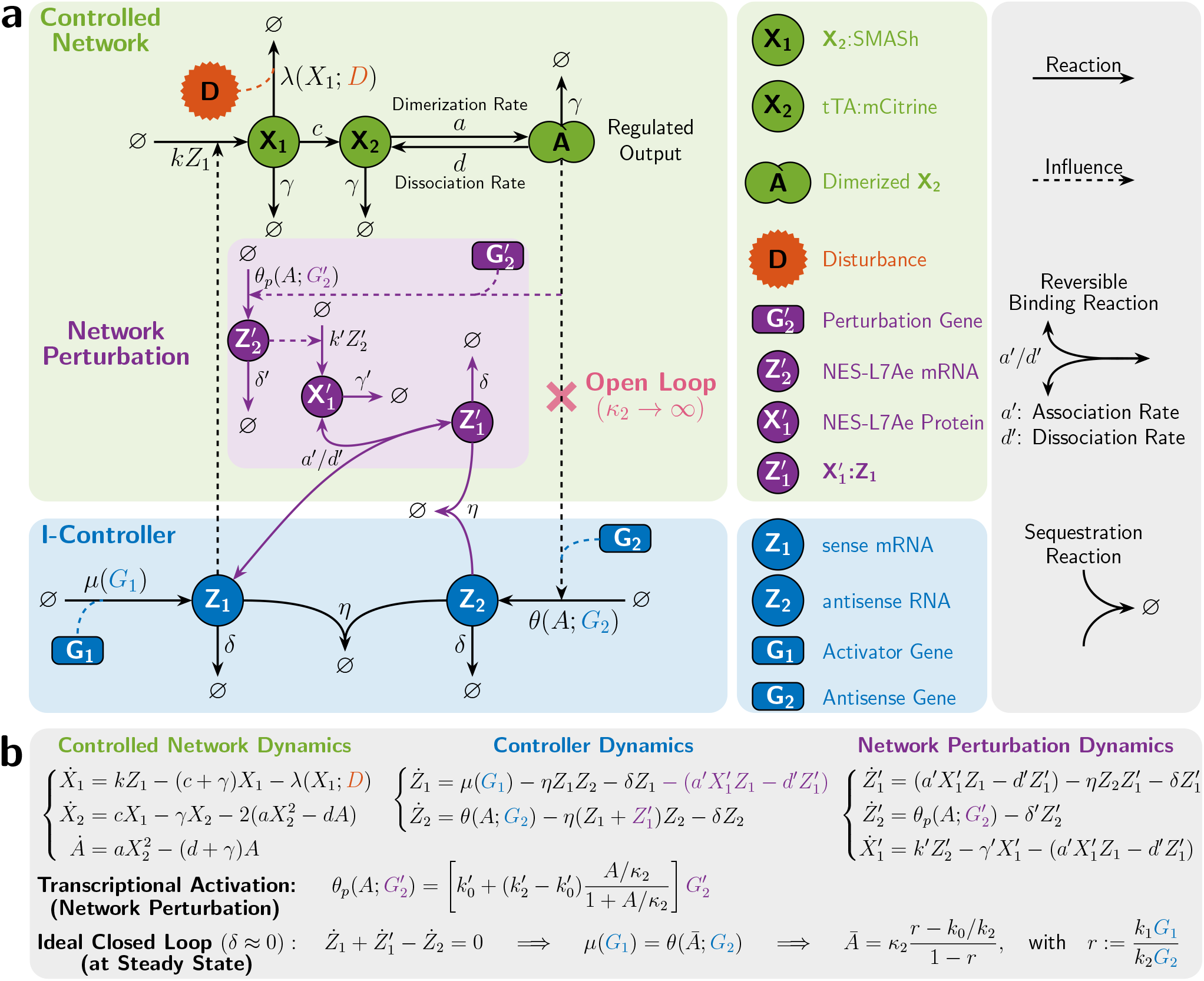
Mathematical Modeling of the I-Circuit with Network Perturbation in Figure 3. **(a)/(b) Schematic/Mathematical Description of the Reduced Model.** This is a special case of the compact model presented in Figure 5(a) where the proportional controller is removed, that is 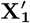 cannot be produced from **Z_1_**. As a result, this network models the integral control action and the network perturbation introduced by the NES-L7Ae gene, denoted by 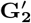 which gives rise to the sub-network in purple. Here, the dimer **A** acts as a transcription factor for both genes **G_2_** and 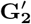. When 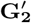 is activated, it is transcribed into 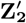 at a rate 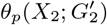 which in turn is translated into 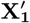 at a rate 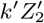. Then 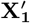 is capable of inhibiting the translation of **Z_1_** by binding to it. Note that the complex 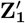 formed from the binding reaction can still sequester the antisense RNA **Z_2_**. In the ideal operation of the antithetic integral controller, where the dilution rate *δ* is negligible with respect to the other rates of the controller, the regulated output **A** has a steady-state concentration, denoted by *Ā*, that is unaffected by the network perturbation. This ensures robust perfect adaptation of the output not only to external disturbances in the controlled network as illustrated in Figure 3, but also to network perturbations as well.

One additional conservation law is appended here to the previous conservation laws in (2). It is given in terms of the total concentration of the bound and free NES-L7Ae gene denoted by 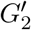. That is, we have

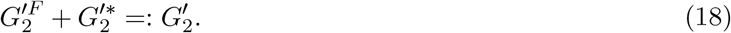

Note that 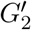 is a constant that is considered to act as an external perturbation to the circuit. Since the binding reactions are much faster than the other reactions in the network (Assumption 1), one can invoke the Quasi-Steady-State Approximation (QSSA) as follows

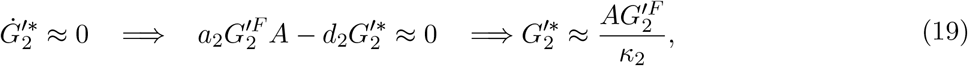

where the dissociation constant *κ*_2_ is given in Table D.13 and is assumed to be equal to that in Table C.9 since **G_2_** and 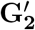 have the same promoters. By substituting the quasi-steady-state approximation of 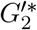 in the conservation law 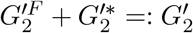, we obtain the following expressions

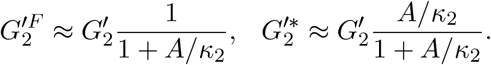

Equipped with these quasi-steady-state approximations, we can update the ODEs of *Z*_1_ and *Z*_2_ from (4) and write down the additional ODEs for 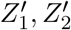 and 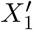.

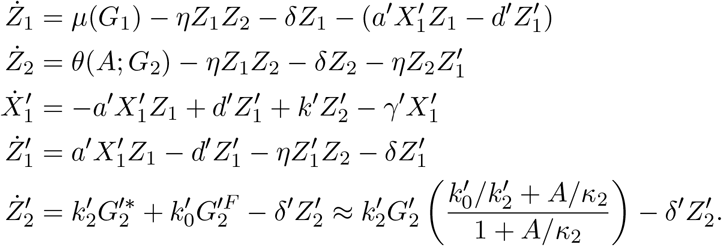

Finally, the dynamics of the reduced model can be written as

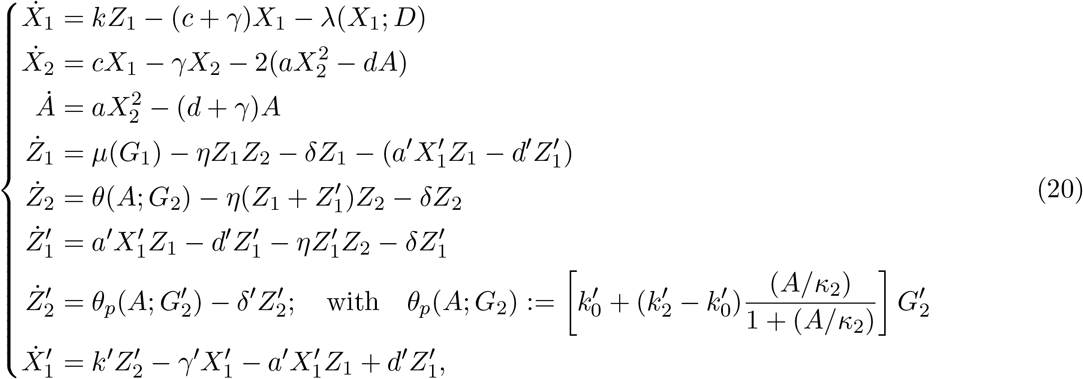

where 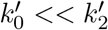 (since leaky transcription is usually much slower than activated transcription) and *λ, μ, θ* are all functions given in (4). Note that by setting 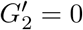, we obtain the circuit in Figure C.7(a) where there is no network perturbation.

### D.3 Model Calibration to the Experimental Data

In this section, we calibrate the model depicted in Figures D.9(a) and (b) to fit the experimentally collected data at steady state. The mathematical model of the measurement described in Supplementary Information C.4 is applicable here as well. However we add here the argument 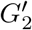 to the measurement function 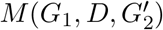 to explicitly show the dependence of the measurement on the concentration of the gene 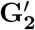. The experimental data are collected for both open- and closed-loops with/without disturbance and with/without network perturbation over two plasmid ratios *G*_1_/*G*_2_ as shown in Figure 5(c). The disturbance is introduced via *D* = 30 nm, while network perturbation is introduced via 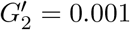 pmol.

#### D.3.1 Choice of Parameter Groups

Consider the circuit depicted in Figures D.9(a) and (b). The measured output for the open- (*i* = *o*) and closed-loop (*i* = *c*) settings are given by

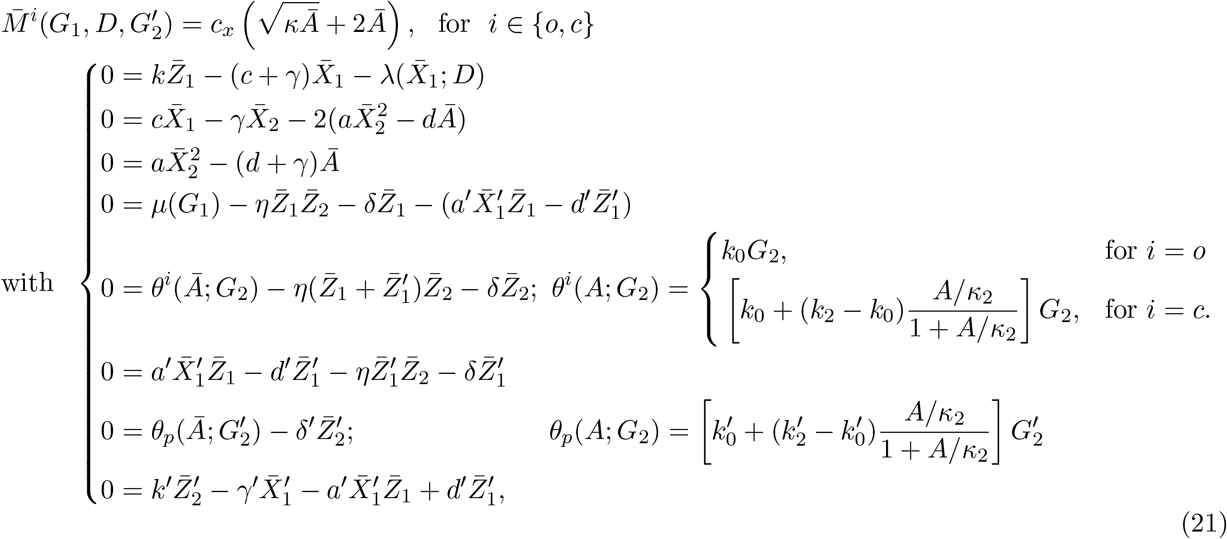

The system model has 7 additional parameters (compared to the circuit in Figure C.7) to be calibrated to the data: 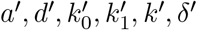 and *γ′*. Once again, we carry out a suitable choice of re-parameterization to obtain a minimal number of (lumped) parameter groups that can be uniquely identified from the steadystate measurements. To specify the choice of the parameter groups, we first express all the variables as rational functions of 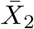. The first three and seventh equations in (21) can be rewritten as

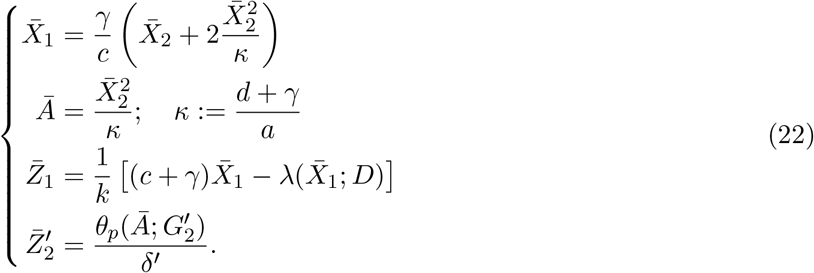

Hence, we expressed now 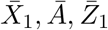 and 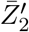 as rational functions of 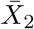. Next, we express 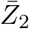 as a rational function of 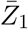 and *Ā* (and thus 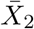). To do so we obtain the following equations

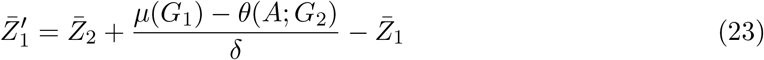

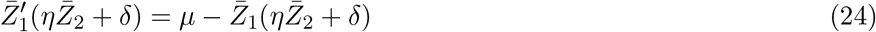

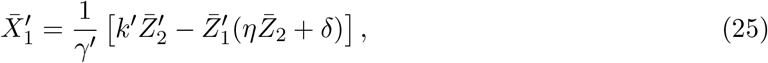

where the first equation is obtained by subtracting the fifth equation in (21) from the sum of the fourth and sixth equations, the second equation is obtained by summing up the fourth and sixth equations in (21), and the third equation is obtained by summing up the sixth and eighth equations in (21). By substituting for 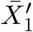 in the sixth equation of (21), we obtain

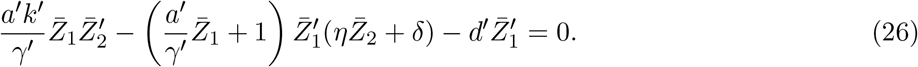

By substituting the expressions for 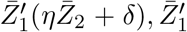 and 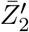 from (24), (23) and (22), respectively we obtain

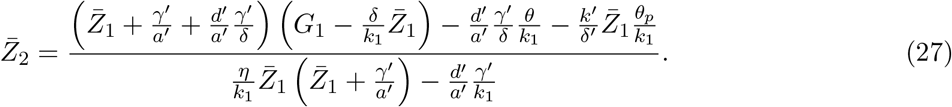

Finally, one can substitute the expressions for 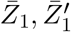 and 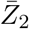 in the fifth equation of (21) to obtain a single (high order) polynomial equation in 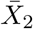 solely which can be solved efficiently in Matlab using the command “root”, instead of solving the set of nonlinear algebraic equations (21).

Next, we rewrite the obtained equations in terms of the parameter groups and transformed variables. By recalling the transformed variable 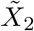 given in (15) and the parameter groups given in (14) and introducing the following five additional parameter groups

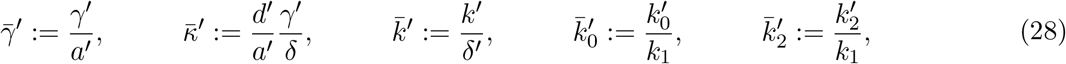

we can rewrite 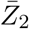 as

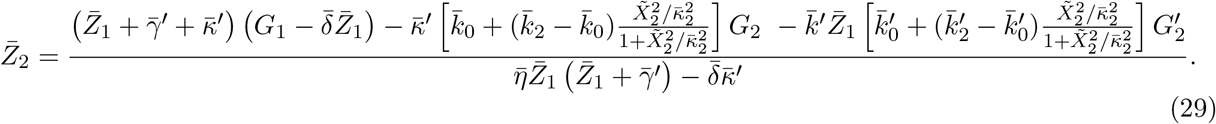

Therefore, the steady-state measurements can be rewritten in terms of the parameter groups and transformed variables as

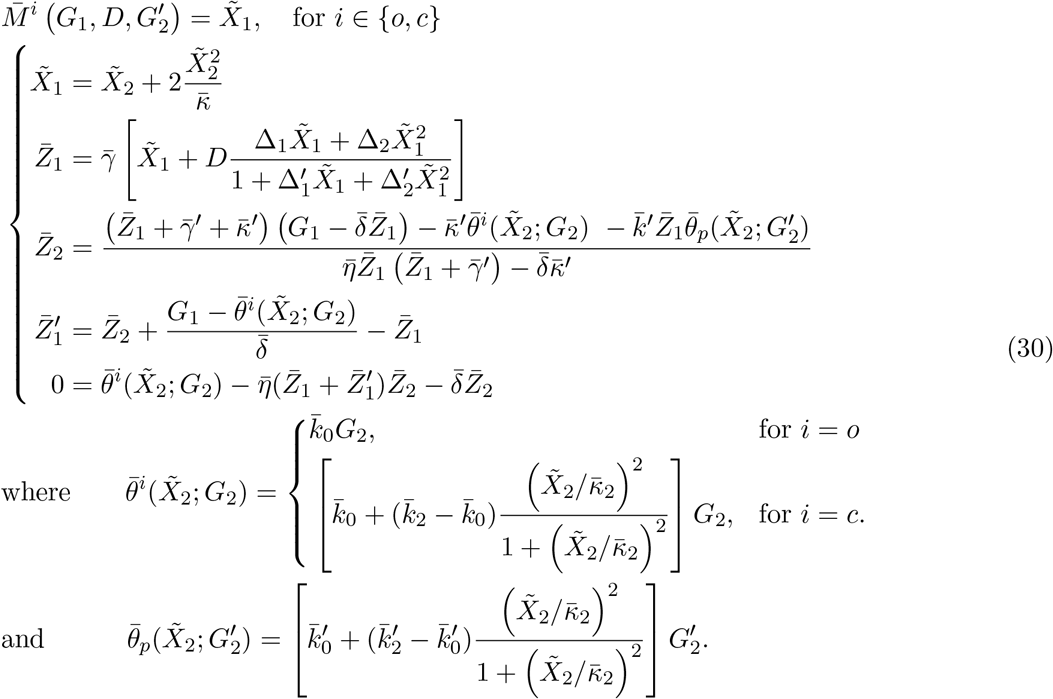

Note that *i* = *o* and *i* = *c* correspond to the open- and closed-loop settings, respectively. Hence given the input *G*_1_, the perturbation 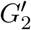, and the disturbance *D*, one can use (30) to compute the mCitrine measurement in the open- and closed-loop settings. We close this section by observing that we have now reduced the parameters to be calibrated down to 16 (as compared to 23).

#### D.3.2 Model Calibration Steps

The model fitting is carried out in three steps to avoid over-fitting. In the first step, the model without network perturbation 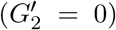 that is obtained in Figure C.7 via the estimated group parameters given in (17) is re-calibrated to the new experimental conditions such as the change in the fluorescence proportionality constant *c_x_*. In the second step, the parameters 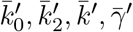 and 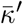 are estimated using the data of the scenario with network perturbation (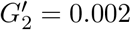 pmol) but without disturbance (*D* = 0 nm). Finally, a prediction step is carried out to further assess the model fitting procedure. In this step, the estimated parameters are used to predict the scenario where both network perturbation and disturbance are applied simultaneously.

##### Estimated Parameter Groups

The model fit and prediction are shown in Figure 5(c), where the optimally estimated parameter groups are given by

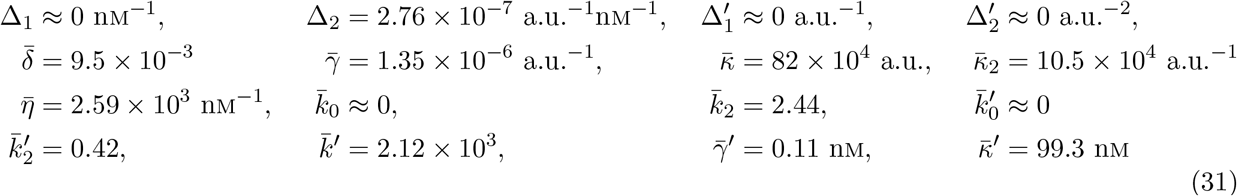

It is straight forward to see that the parameter groups that are common with (17) are numerically very close if one takes into consideration that the fluorescence proportionality constant *c_x_* is increased by seven to eight times.

## E Mathematical Modeling of the Circuit in Figure 4

Consider the circuit depicted in Figure 4(a), with a Proportional-Integral controller, that can operate in either open or closed loop. We first present a detailed (mechanistic) mathematical model and then carry out a model reduction technique that allows us to analyze the steady-state behavior. Finally we provide the technical details of fitting the model to the experimentally obtained data.

### E.1 Full Model Description

A detailed biochemical reaction network that models the dynamics of the circuit in Figure 4(a) can be obtained by replacing the translation reaction in Table C.9 with a slightly modified version, and appending additional inhibition, degradation/dilution and sequestration reactions depicted in Tables E.14 and E.15. These species and reactions describe the biochemical reaction sub-network that provides a proportional control action via the additional protein 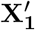 which is translated from the same mRNA **Z_1_** as **X_1_**. This protein can bind to **Z_1_** to form the complex 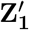 to inhibit translation. However, the complex 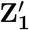 can still be sequestered by the anti-sense RNA **Z_2_**.

**Table E.14:** List of Additional Biochemical Species (PI-Circuit)

| Species | Symbol | Species | Symbol |
| --- | --- | --- | --- |
| NES-L7Ae | $\mathbf{X}'_1$ | $\mathbf{Z}_1:\mathbf{X}'_1$ | $\mathbf{Z}'_1$ |

### E.2 Model Reduction

In this section, the full model given in Tables C.9 and E.15 is mathematically reduced to the model described schematically in Figure E.10(a) and mathematically in E.10(b). The model reduction procedure is based on the reduced model that is obtained previously in Supplementary Information C.2. In fact, one can update the ODEs of *Z*_1_ and *Z*_2_ from (4) and write down the additional ODEs for 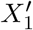 and 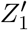.

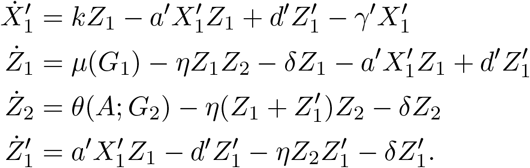

**Table E.15:**
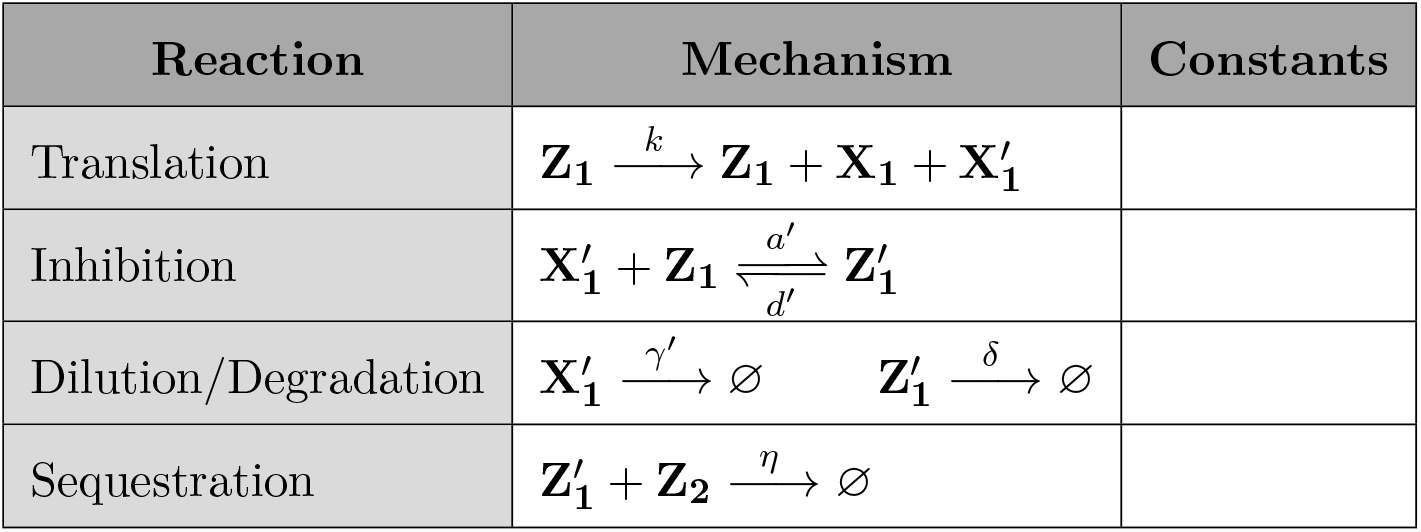
List of Additional Biochemical Reactions (Pl-Circuit)

Finally, the dynamics of the reduced model is thus given by

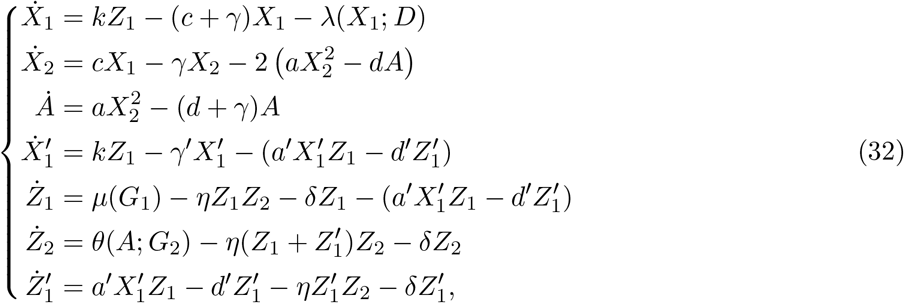

where *λ, μ* and *θ* are all functions given in (4).

### E.3 Model Calibration to the Experimental Data

In this section, we calibrate the model depicted in Figures E.10(a) and (b) to fit the experimentally collected data at steady state. The mathematical model of the measurement described in Supplementary Information C.4 is applicable here as well. The experimental data are collected for both open- and closed-loops with/without disturbance and with/without Proportional P-control over three different plasmid ratios *G*_1_/*G*_2_ as shown in Figure 5(d). The disturbance is introduced via *D* = 30 nm.

**Figure E.10:**
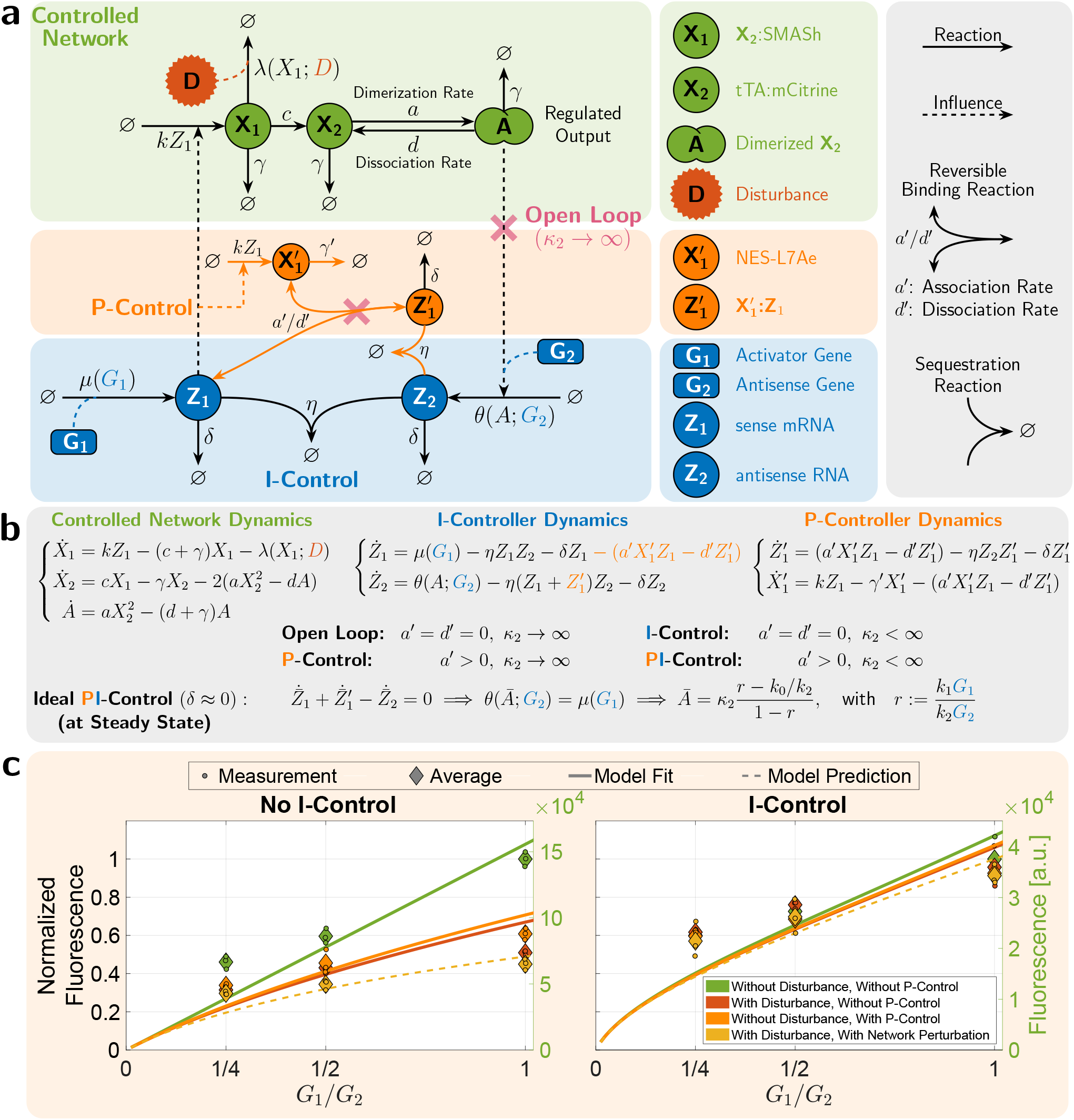
Mathematical Modeling of the PI-Circuit in Figure 4. **(a)/(b) Schematic/Mathematical Description of the Reduced Model.** This is a special case of the compact model presented in Figure 5(a) where the network perturbation is removed, that is the additional gene 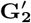 is removed. As a result, this network models the Proportional-Integral (PI) control actions. The P-controller, depicted in the orange box, is realized via the production of 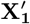 by **Z_1_** at the same rate *k* as that of **X_1_**. This allows 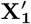 to act as a proxy for **X_1_**. A negative feedback action is then achieved via the (un)binding reaction between 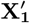 and **Z_1_** which inhibits the production of **X_1_**. In the ideal operation of the controller, where the dilution rate *δ* is negligible with respect to the other rates of the controller, the regulated output **A** has a steady-state concentration, denoted by *Ā*, that is unaffected by the P-controller. This ensures that the robust perfect adaptation of the regulated output is not influenced by the proportional controller. In fact, the P-controller has the effect of shaping the transient dynamics and reducing the steady state variance of **A** while leaving *Ā* unchanged.

#### E.3.1 Choice of Parameter Groups

Consider the circuit depicted in Figures E.10(a) and (b). The measured output for the open- (*i* = *o*) and closed-loop (*i* = *c*) settings are given by

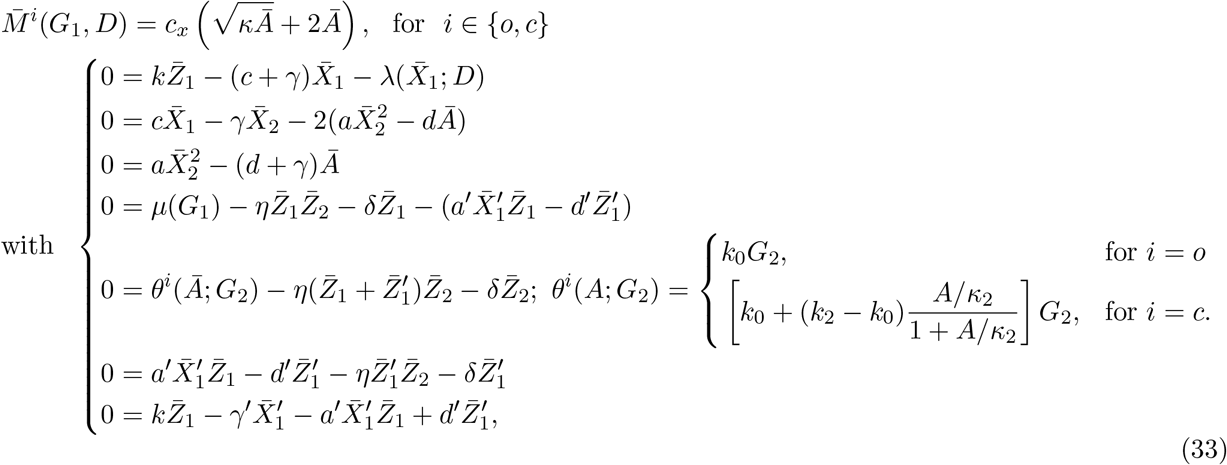

Note that by open loop (resp. closed loop), we mean the circuit without (resp. with) the integral controller. The system model has 3 additional parameters (compared to the circuit in Figure C.7) to be calibrated to the data: *a′, d′* and *γ′*. Once again, we carry out a suitable choice of re-parameterization to obtain a minimal number of (lumped) parameter groups that can be uniquely identified from the steadystate measurements. To specify the choice of the parameter groups, we first express all the variables as rational functions of 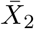. The first three equations in (33) can be rewritten as

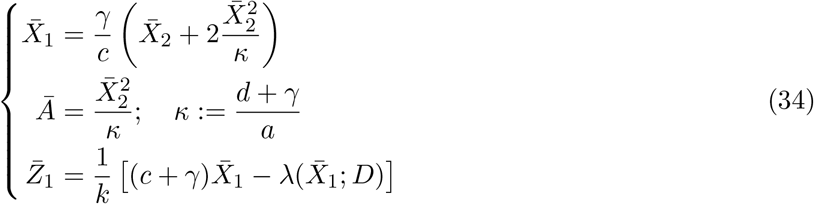

Hence, we expressed now 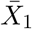, *Ā* and 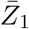 as rational functions of 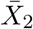. Next, we express 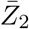 as a rational function of 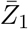 and *Ā* (and thus 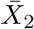). To do so we obtain the following equations

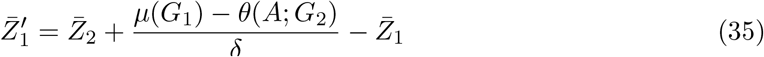

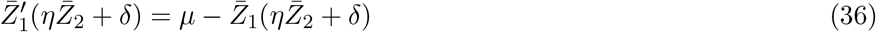

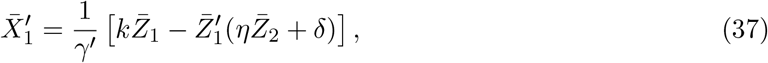

where the first equation is obtained by subtracting the fifth equation in (33) from the sum of the fourth and sixth equations, the second equation is obtained by summing up the fourth and sixth equations in (33), and the third equation is obtained by summing up the last two equations in (33). By substituting for 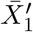 in the sixth equation of (33), we obtain

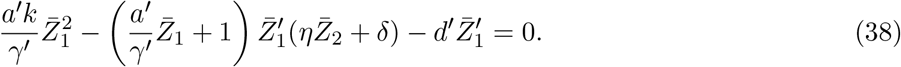

By substituting the expressions for 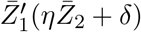 and 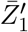 from (36) and (35), respectively we obtain

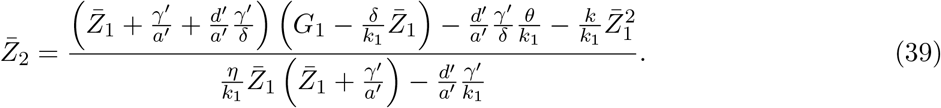

Finally, one can substitute the expressions for 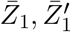 and 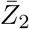 in the fifth equation of (33) to obtain a single (high order) polynomial equation in 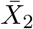 solely which can be solved efficiently in Matlab using the command “root”, instead of solving the set of nonlinear algebraic equations (33).

Next, we rewrite the obtained equations in terms of the parameter groups and transformed variables. By recalling the transformed variable 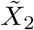 given in (15) and the parameter groups given in (14) and introducing the following three additional parameter groups

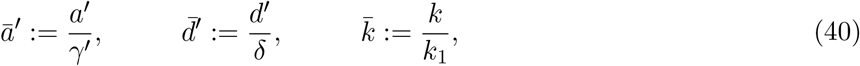

we can rewrite 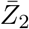 as

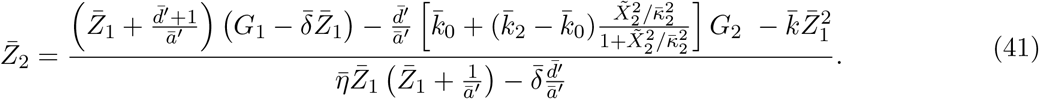

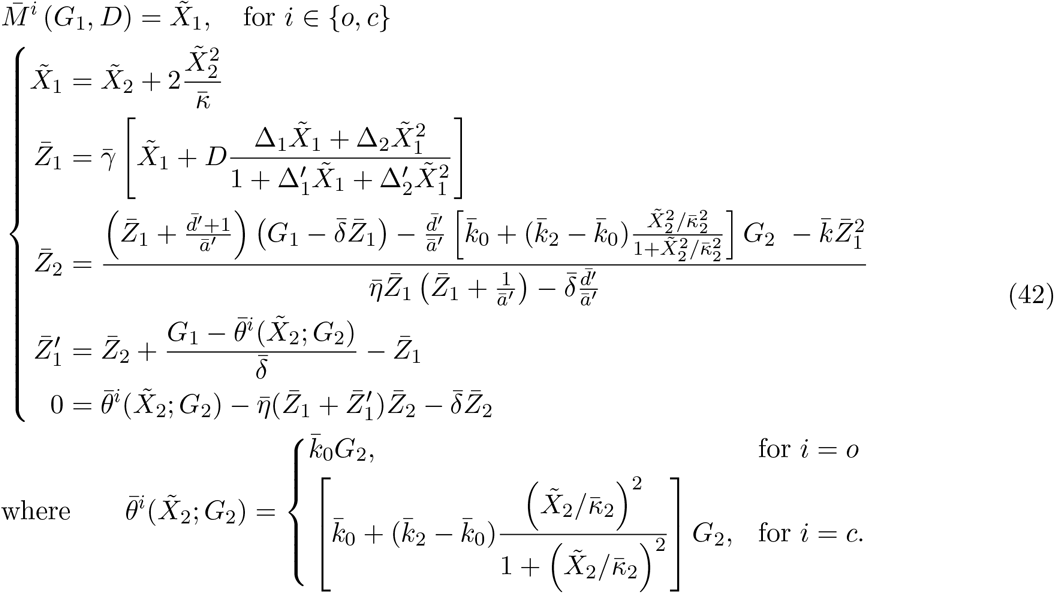

Recall that *i* = *o* and *i* = *c* correspond to the open- and closed-loop settings, respectively. Hence given the input *G*_1_ and the disturbance *D*, one can use (42) to compute the mCitrine measurement in the open- and closed-loop settings. We close this section by observing that we have now reduced the parameters to be calibrated down to 14 (as compared to 19).

#### E.3.2 Model Calibration Steps

The model fitting is carried out in three steps to avoid over-fitting. In the first step, the model without a proportional controller (the production of 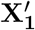 by **Z_1_** is removed) that is obtained in Figure C.7 via the estimated group parameters given in (17) is re-calibrated to the new experimental conditions such as the change in the fluorescence proportionality constant *c_x_*. In the second step, the parameters 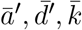 are estimated using the data of the scenario with a proportional controller but without disturbance (*D* = 0 nm). Finally, a prediction step is carried out to further assess the model fitting procedure. In this step, the estimated parameters are used to predict the scenario where both a proportional controller and disturbance are applied simultaneously.

##### Estimated Parameter Groups

The model fit and prediction are shown in Figure 5(d), where the optimally estimated parameter groups are given by

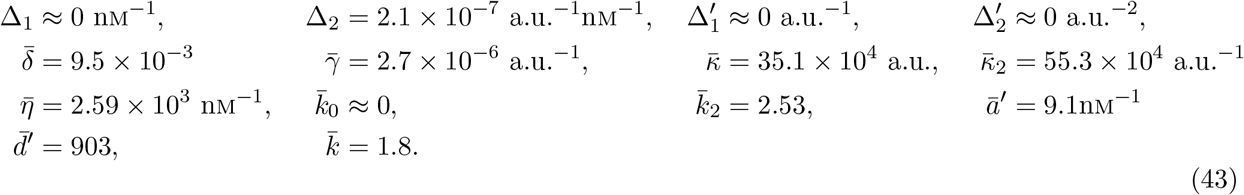

## F Antithetic Proportional-Integral Control of Plasma Glucose

In this section, we provide the mathematical details underlying the simulation results given in Figure 7 which demonstrate the tight regulation of glucose concentration in the plasma via an antithetic proportional-integral controller.

### F.1 Brief Description of the Glucose-Insulin Network to be Controlled

The mathematical whole-body model describing the dynamics of the glucose-insulin system is adopted from [38]. The FDA-approved mathematical model is comprised of 15 species, 19 reactions and 64 parameters and is conveniently implemented as part of the SimBiology toolbox in MATLAB [49]. It captures the dynamics of glucose and insulin across the various relevant organs/tissues in the body. For a healthy subject, the model parameters take particular values; whereas for a type-II diabetic patient, certain relevant parameters are modified to reflect the lower sensitivity to insulin (see [38, Table I] and [49] for details). In contrast, for a type-I diabetic patient, the endogenous insulin production reactions [38, Equations 23–26] are removed from the model to reflect the death or inactivity of the *β*-cells while the parameters are kept the same as those associated with a healthy subject. The output to be controlled here is the plasma glucose concentration in mg/dL; whereas the actuated input is the total secreted quantity of insulin in the plasma in pmol.

### F.2 P-Type Proportional-Integral Control Motif

In the previous circuits of Figures 2, 3, 4 and 6, the controlled networks have positive gains. That is, producing more input species leads to an increase in the output species. As a result, the designed controllers for these networks implement negative feedback to ensure closed-loop stability and are hence called N-type controllers (for Negative feedback). In contrast, the glucose-insulin network to be controlled here has a negative gain since producing more insulin (input species) leads to a decrease in plasma glucose levels (output species). Subsequently, the controller for this network should implement positive feedback (P-type) to achieve overall negative feedback for the closed loop. A P-type antithetic integral controller can be achieved by switching the roles of the sense and antisense RNAs, and a P-type proportional controller can be achieved by using a promoter that is activated in the presence of glucose to drive the expression of insulin.

Next, we provide the controller differential equations that we append to the glucose-insulin model. Let *G* and *I* denote the plasma glucose concentration (output) and total plasma insulin molecules (input) in mg/dL and pmol, respectively. The controller dynamics are thus given by

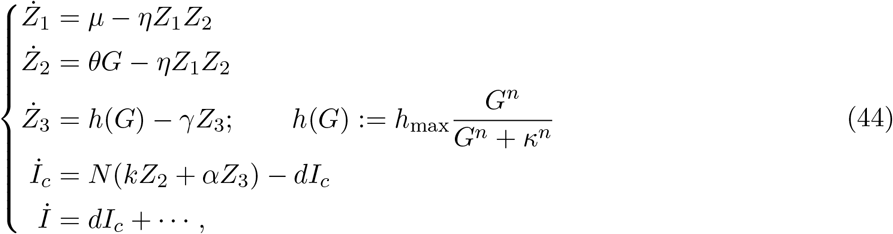

where *Z*_1_ and *Z*_2_ represent the average quantities (per cell) of the anti-sense and sense RNAs, respectively, while *Z*_3_ represents the average quantity of another mRNA that is transcribed by the gene associated with the proportional controller (see Figure 7(a)). Note that the average quantities are taken across *N* cells in pmol. Furthermore, *I_c_* denotes the total quantity of produced insulin in all the cells before they diffuse at a rate *d* to the plasma. Hence the rate of total insulin secretion into the plasma given by *dI_c_* (in pmol/h) serves as the proportional-integral actuation to the glucose-insulin system. Note that the controller reactions and parameters are appended to the SimBiology glucose-insulin model in Matlab to close the loop.

### F.3 Choice of Controller Parameter Values

To enhance the performance of any controller, the control parameters has to be properly tuned. However, in practice, the various biological controller parameters cannot be freely tuned since the time scales are governed by gene expression processes. The controller parameter values appearing in the system of differential equations of the controller (44) are listed in Table F.16 for Type I and II diabetic subjects.

**Table F.16:** List of plausible numerical values of the various biological controller parameters for controlling both Type I and Type II diabetic patients by the Proportional-Integral controller described in (44). Standalone proportional control is achieved by setting *k* = 0, while standalone integral control is achieved by setting *α* = 0.

| Controller Parameters | Numerical Values |  | Units | Brief Description |
| --- | --- | --- | --- | --- |
|  | Type I | Type II |  |  |
| $\mu$ | $1.66 \times 10^{-10}$ | $1.66 \times 10^{-10}$ | pmol/h | Constitutive transcription rate |
| $\theta$ | $\mu/100$ | $\mu/100$ | pmol dL/mg/h | Transcription rate constant |
| $\eta$ | $10^{10}$ | $10^{10}$ | 1/pmol/h | Transcription rate |
| $N$ | $10^9$ | $10^9$ | Dimensionless | Number of implanted controller cells |
| $k$ | $10N$ | $10N$ | 1/h | Translation rate constant |
| $\alpha$ | $700N$ | $700N$ | 1/h | Translation rate constant |
| $\delta$ | $\log(2)/0.5$ | $\log(2)/0.5$ | 1/h | Fast mRNA degradation rate constant |
| $d$ | $1/0.5$ | $1/0.5$ | 1/h | Insulin diffusion rate constant |
| $h_{\max}$ | $\mu$ | $2.5\mu$ | pmol/h | Transcription rate |
| $\kappa$ | 200 | 150 | mg/dL | Dissociation constant |
| $n$ | 4 | 4 | Dimensionless | Hill coefficient |

Next, we provide the rationale behind picking realistic numerical values of the various biological control parameters in the case of Type I diabetic subjects. This ensures that the modeling and simulation study summarized in Figure 7 is numerically realistic and plausible. In the case of Type II diabetic subjects, the control parameters are similar to the Type I case, with some additional fine tuning to enhance the performance. Two important numbers are of particular interest: transcription and translation rates. It is shown in [24] that the transcription rate ranges between 0.1 and 100 mRNAs per hour with extreme cases going up to more than 500 mRNAs per hour. Hence picking a transcription rate, for the antisense RNA **Z_1_**, of 100 mRNAs per hour allows us to set *μ* to

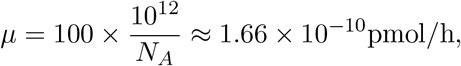

where *N_A_* is Avogadro’s number. Since *h*(*G*) is also a transcription rate whose maximum value (as *G* → ∞) is *h*_max_, then we also set

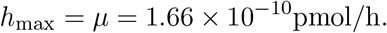

The desired set-point of the controlled glucose levels in the plasma is chosen here to be 100mg/dL. Hence for the proportional-integral controller to achieve this set-point given by *μ/θ*, we set *θ* to be

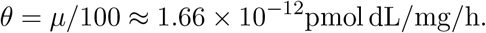

Observe that for this value of *θ*, the transcription rate of the sense-mRNA (around the set-point) is given by *μ* which is already selected to be plausible. Next, it is also shown in [24] that the translation rate ranges between 1 and 1000 proteins per mRNA per hour. This allows us to set *k* and *α* to plausible values between 1 and 1000 h^−1^ multiplied by the number of implanted cells N which is taken to be 1 billion — equal to the number of *β*-cells in the human pancreas [50]. The sequestration rate *η* is chosen to be large enough to reflect a fast sense/antisense RNA hybridization. Furthermore, *κ* is selected to tune the threshold of the Hill function with high sensitivity (Hill coefficient *n* = 4).

Ideally, a proportional controller exhibits an instantaneous feedback from the output into the input. Here, in a more practical setting, the proportional controller can be realized via gene expression (see Figure 7(a)) where the degradation rate of the associated mRNA **Z_3_** is desired to be fast to mimic the ideal instantaneous proportional control action. Consequently, the degradation rate of the mRNA **Z_3_** is picked to be fast enough, that is log(2)/0.5h^−1^ to reflect a relatively short half-life of 0.5 h. Finally, a conversion reaction 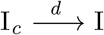 is introduced to reflect a delay caused by the diffusion of the insulin from the cells to the plasma at a rate of *d* = 1/0.5h^−1^. This reflects a 30 min average time to secrete 1000 newly produced insulin molecules into the plasma.

